# Oxytocin Acts on Astrocytes in the Central Amygdala to Promote a Positive Emotional State

**DOI:** 10.1101/2020.02.25.963884

**Authors:** Jérôme Wahis, Damien Kerspern, Ferdinand Althammer, Angel Baudon, Stéphanie Goyon, Daisuke Hagiwara, Arthur Lefèvre, Benjamin Boury-Jamot, Benjamin Bellanger, Marios Abatis, Miriam Silva da Gouveia, Diego Benusiglio, Marina Eliava, Andrej Rozov, Ivan Weinsanto, Hanna Sophie Knobloch-Bollmann, Hong Wang, Marie Pertin, Perrine Inquimbert, Claudia Pitzer, Jan Siemens, Yannick Goumon, Benjamin Boutrel, Pascal Darbon, Christophe Maurice Lamy, Javier E. Stern, Isabelle Décosterd, Jean-Yves Chatton, W. Scott Young, Ron Stoop, Pierrick Poisbeau, Valery Grinevich, Alexandre Charlet

**Affiliations:** Centre National de la Recherche Scientifique and University of Strasbourg, UPR3212 Institute of Cellular and Integrative Neurosciences, Strasbourg, France; German Cancer Research Center (DKFZ), Heidelberg, Germany; Department of Neuropeptide Research for Psychiatry, Central Institute of Mental Health, University of Heidelberg, Mannheim, Germany; Center for Psychiatric Neurosciences, Hôpital de Cery, Lausanne University Hospital (CHUV), Lausanne, Switzerland; OpenLab of Neurobiology, Kazan Federal University, Kazan, Russia and Department of Physiology and Pathophysiology, University of Heidelberg, Heidelberg, Germany; Department of Molecular and Cellular Biology, Center for Brain Science, Harvard University, Cambridge, USA; Department of Pharmacology, Heidelberg University, Heidelberg, Germany; Pain center, Department of Anesthesiology, Lausanne University Hospital (CHUV), Lausanne, Switzerland; Interdisciplinary Neurobehavioral Core (INBC), Ruprecht-Karls-Universität, Heidelberg; Division of Anatomy, Faculty of Medicine, University of Geneva, Geneva, Switzerland; Center for Neuroinflammation and Cardiometabolic Research, Georgia State University, Atlanta, USA; Department of Fundamental Neurosciences, Faculty of Biology and Medicine (FBM), University of Lausanne, Lausanne, Switzerland; Section on Neural Gene Expression, National Institute of Mental Health, National Institutes of Health, Bethesda, MD, USA; University of Strasbourg Institute for Advanced Study (USIAS), Strasbourg, France

## Abstract

Oxytocin orchestrates social and emotional behaviors through modulation of neural circuits in brain structures such as the central amygdala (CeA). The long-standing dogma is that oxytocin signaling in the central nervous system occurs exclusively via direct actions on neurons. However, several findings over the last decades showed that astrocytes actively participate in the modulation of neuronal circuits. Here, we investigate the degree of astrocytes’ involvement in oxytocin functions. Using astrocyte’ specific gain and loss of function approaches, we demonstrate that CeA astrocytes not only directly respond to oxytocin, but are actually necessary for its effects on neuronal circuits and ultimately behavior. Our work identifies astrocytes as a crucial cellular substrate underlying the promotion of a positive emotional state by oxytocin. These results further corroborate that astrocytes are key regulators of neuronal circuits activity by responding to specific neuropeptidergic inputs, and opens up new perspectives to understand how neuromodulators gate brain functions.

## INTRODUCTION

The neuropeptide oxytocin (OT) modulates key neurophysiological functions ^1^, including anxiety and pain, notably through its action on central amygdala (CeA) microcircuits^2–5^. Oxytocin receptors (OTR) are expressed in the lateral and capsular part of the CeA (CeL)^3^. In this nucleus, OTR activation leads to increased firing of γ-aminobutyric acid (GABA) expressing interneurons. This directly inhibits projection neurons of the medial part of the CeA (CeM), the functional output of the CeA, thereby leading to clear behavioral outputs^3, 4, 6^. Besides, the CeA is recognized as a major player in the pathophysiology of several neurological diseases, among which is neuropathic pain^7^, for which recent studies also suggest a critical involvement of astrocytes^8–10^ as well as the oxytocinergic system^11, 12^. However the possible involvement of astrocytes in the neuromodulatory effects of oxytocin has rarely been explored^13–16^. That, despite the numerous findings of astrocytes active involvement in the regulation of neural circuits in a brain area specific manner^17–20^, notably in the CeA^21^, with yet many controversies remaining about the mechanisms involved^22, 23^.

Herein, we employed patch-clamp techniques, calcium imaging, and behavioral assays to measure the consequences of *in vivo* and *ex vivo* direct manipulation of CeL astrocytes in both rats and mice. To this end, we used pharmacological approaches as well as genetic tools to ablate OTRs or expressed opsins specifically in CeL astrocytes. We demonstrate that astrocytes are a necessary component in the oxytocinergic modulation of CeA neuronal circuits and propose a mechanism by which astrocytes relay OT effects in the CeA by gating neuronal N-methyl-D-aspartate receptors (NMDAR) activation through release of the co-agonist D-serine. This ultimately leads to promotion of the behavioral correlates of positive emotional states. With this study, we show for the first time that astrocyte-neuron communication is crucial for an efficient integration of positive emotions within the CeA, a structure involved in the processing of emotionally-relevant cues.

## RESULTS

### CeL Astrocytes Express Functional Oxytocin Receptors

To specifically investigate whether CeA astrocytes express OTRs, we performed fluorescence *in situ* hybridizations (FISH) on rat CeA sections and found overlap between OTR mRNA and an astrocyte marker, glutamine synthetase (Figure 1a-c). We quantified cells positive for both markers and found that a randomly (Figure S1a) distributed sub-population of about 13.6% of CeL astrocytes expresses the OTR mRNA (Figure 1c). We further corroborated this finding using the aldehyde dehydrogenase 1 family member L1 (ALDH1L1) as an astrocyte marker (Figure S1a), and confirmed it to be comparable in mice CeL astrocytes (Figures S1c-d; 2c).

**Figure 1.**
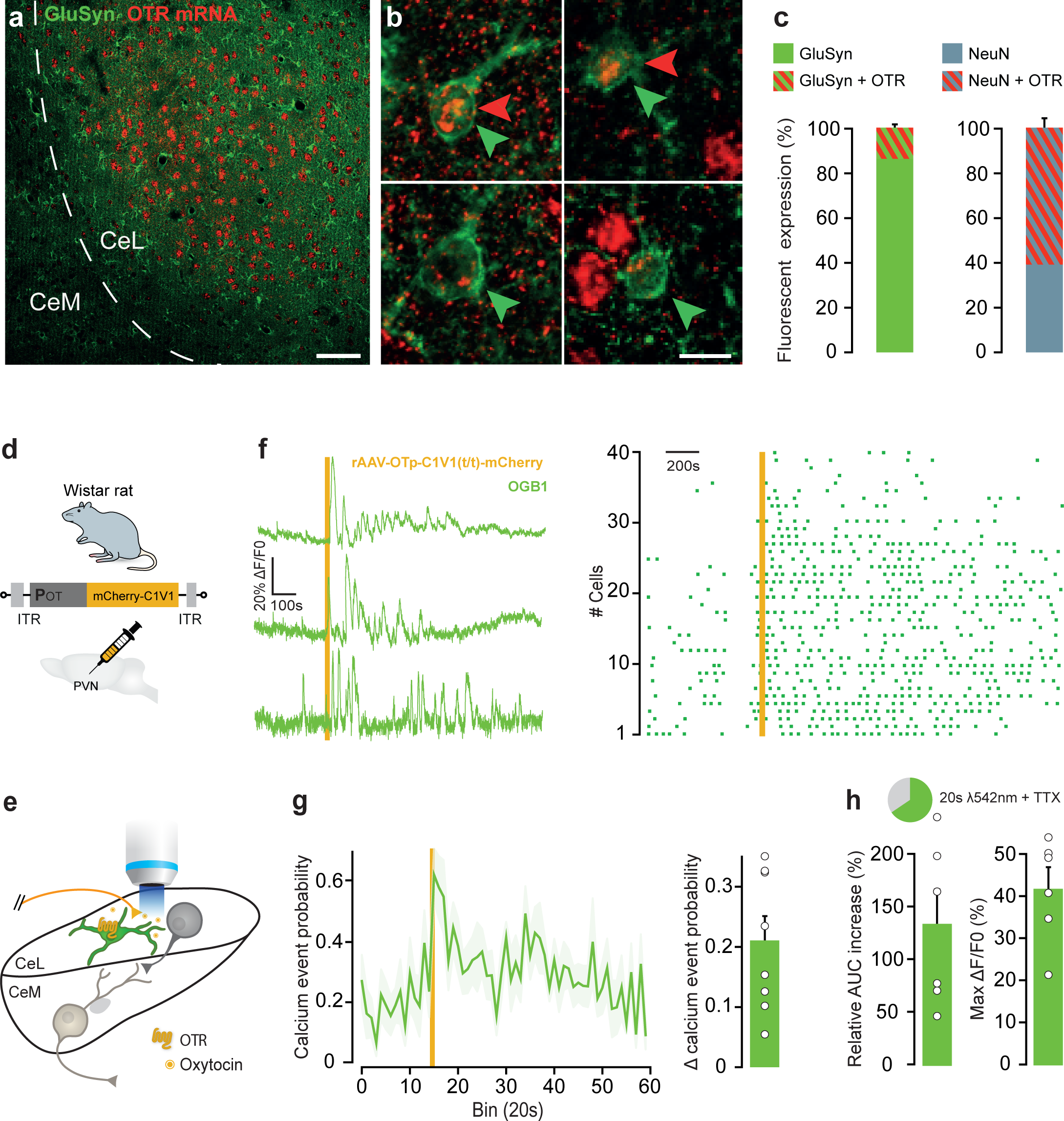
CeL Astrocytes Express Functional Oxytocin Receptors. (**a**) Overview of CeA fluorescent *in situ* hybridization of OTR mRNA (red) and glutamine synthetase immunostaining (GluSyn, green). (**b**) High magnification images of cells positive for OTR mRNA and/or GluSyn (double arrows); green arrows point GluSyn positive cells; red arrows point OTR mRNA-positive cells. Scale bars 100 (a) and 10µm (b). (**c**) Proportion of CeL astrocytes (GluSyn positive cells, left) and neurons (NeuN positive cells, right) positive for OTR mRNA (red, hatched) compared to the total number of OTR mRNA positive cells (n astrocytes = 1185, n neurons = 1254, n rats = 3). (**d**) Experimental rat model for acute brain slice calcium imaging. (**e**) Experimental scheme of the horizontal CeA slice preparation used, showing C1V1(t/t) expressing OT axons (yellow) arising from PVN and projecting to the CeL. (**f**-**h**) Effect of the activation of OT axons in the CeL through λ542 nm light pulses (10 ms width, 30 Hz, duration 20 s) on calcium transients of CeL astrocytes. (**f**) (Left) Typical ΔF/F0 traces; (right) raster plot displaying calcium events per responding astrocyte. (**g**) (Left) Time course of the calcium events probability per bin of 20s; (right) Δ calcium events probability before and after light-driven activation of CeL OT axons. (**h**) (Top) Pie chart of the proportion of responsive astrocytes; (bottom) Relative increase of ΔF/F0 AUC over baseline after light-driven activation of CeL OT axons and maximal peak values reached for responsive astrocytes. Data are expressed as means across slices plus SEM (n slices (ns) = 6, n astrocytes (na) = 37). White circles indicate averages across responding astrocytes per slices. (Statistics in Table S1).

To test whether endogenous activation of astrocytic OTR elicits physiological responses in CeL astrocytes, we performed calcium imaging in brain slices of rats using the calcium indicator Oregon Green® 488 BAPTA-1 (OGB1) and identified astrocytes through sulforhodamine 101 labelling (SR101) (Figure S2a). Indeed, measuring changes in intracellular calcium is the most commonly accepted readout of astrocytic activity^24^. We first confirmed that SR101 positive cells of the CeL displayed typical electrophysiological properties of astrocytes (Figure S2b-c). We then expressed the opsin C1V1(t/t) under the oxytocin promoter through rAAVs (rAAV-OTp-C1V1(t/t)-mCherry) injected in the paraventricular, supraoptic, and accessory hypothalamic nuclei of rats, to enable light-evoked activation of oxytocin neurons and their distant axons localized within the CeL^5, 6^ (Figure 1d-e, S1e). We found that light-driven activation of C1V1(t/t) expressing oxytocinergic axons rapidly evoked long-lasting and oscillating calcium transients in astrocytes (Figure 1f-h). These calcium transients persisted during bath application of TTX, thereby excluding the possibility that they were evoked secondary to neuronal circuit activity.

### OTR Activation Evokes Astrocytic Calcium Transients in CeL Astrocyte *Syncitium*

To confirm that the observed increase of intracellular calcium in CeL astrocytes is mediated by direct action on OTRs, we used the exogenous and OTR-selective agonist [Thr4Gly7]-oxytocin (TGOT). TGOT induced calcium transients in astrocytes, which were prevented by the incubation of the OTR antagonist [d(CH2)5,Tyr(Me)2,Orn8]-vasotocin (dOVT) and remained unaffected by the presence of TTX (Figure 2a).

**Figure 2.**
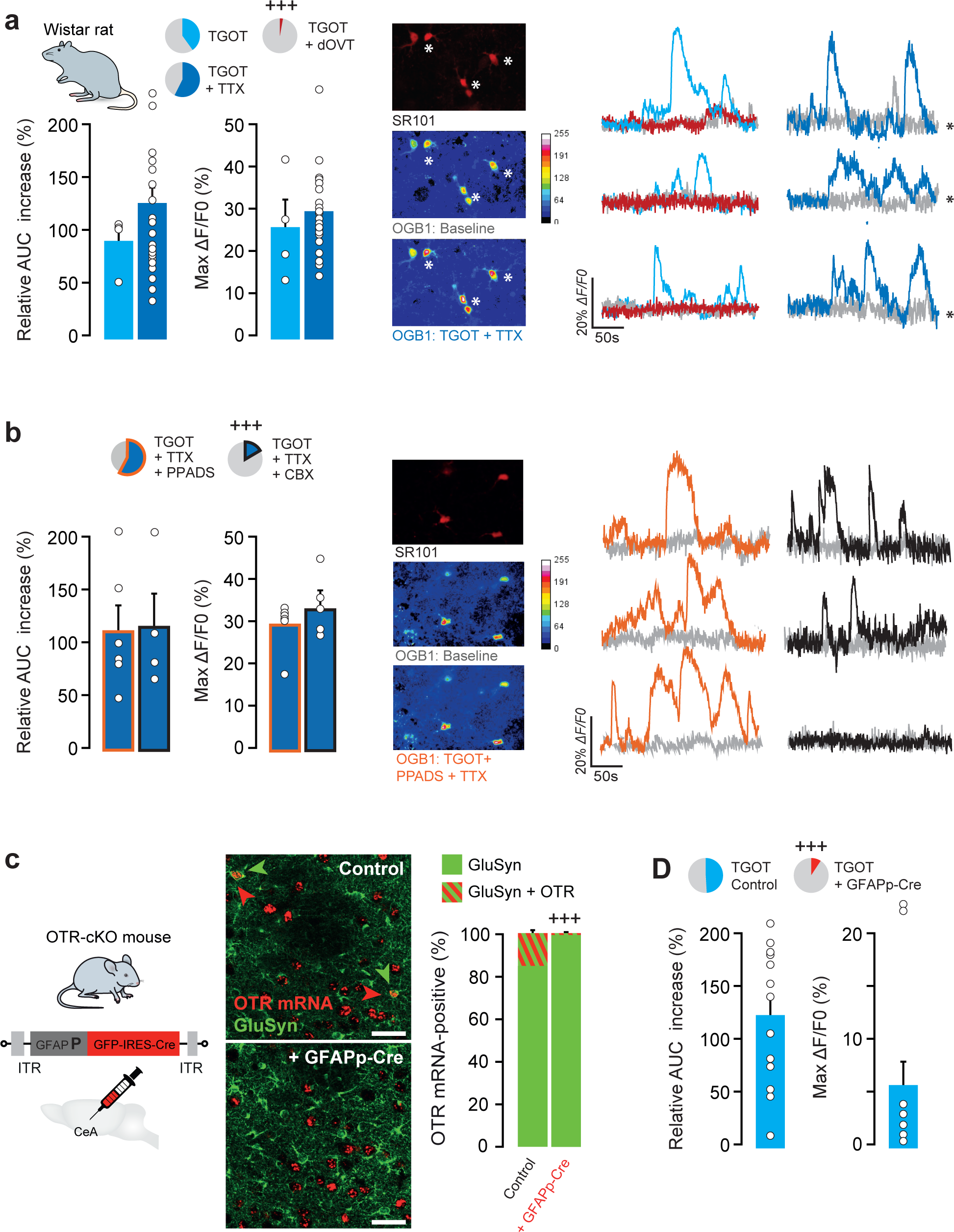
OTR Activation Evokes Astrocytic Calcium Transients in the CeL Astrocyte’ *Syncitium* of Rats and Mice. (**a**-**b**) (Top left) Pie charts display the proportion of responsive astrocytes in rats; (middle) images of CeL astrocytes identified through SR101 (red) and corresponding pseudo-color images of OGB1 fluorescence during baseline and after drug application (stacks of 50 images over 25s of recording). Asterisks indicate astrocytes from which the example ΔF/F0 traces are shown; (right) typical ΔF/F0 traces; (histograms) relative increase of ΔF/F0 AUCs after drug application and maximal peak values reached for responsive astrocytes upon exposition to, from left to right: (**a**) TGOT (400 nM; ns = 4, na = 39; light blue), TGOT+TTX (1 μM; ns = 22, na = 182; dark blue), TGOT+TTX+PPADS (50 μM; ns = 6, na = 59; orange borders) and TGOT+TTX+CBX (100μM; ns=8, na=53; black borders); histograms for dOVT (1 µM; ns = 3, na = 34; red) are not shown, since only 1/34 astrocyte responded. (**c**) Specific deletion of OTRs in mice CeL astrocytes. (Left) Experimental animal model; (middle) example pictures of OTR mRNA (red) and GluSyn labelling in mice injected with rAAV-GFAPp-GFP (top) or rAAV-GFAPp-IRES-Cre vector (bottom); (right) proportion of CeL astrocytes positive for OTR mRNA (red, hatched) compared to total number of GluSyn positive cells (green) (rAAV-GFAPp-GFP vector: n astrocytes = 1340, n mice = 3; rAAV-GFAPp-IRES-Cre vector: n astrocytes = 1561, n mice = 4). (**d**) Histograms displaying relative increase of ΔF/F0 AUCs after drug application and maximal peak values reached for responsive astrocytes in Control vector (rAAV-GFAPp-GFP) injected mice upon exposure to TGOT (400 nM; ns = 13, na = 35; light blue). Histograms for data after Cre induced ablation of CeL astrocytes’ OTRs (ns = 10) are not shown, since only 62/523 astrocytes showed calcium transients modifications. All values were statistically compared to those of TGOT+TTX: ^+++^ P < 0,001, Mann-Whitney U test (Statistics in Table S2).

Next, we sought to reinforce the conclusions drawn from our pharmacological experiments by testing whether the OTRs expressed in CeL astrocytes are responsible for the TGOT-induced increase of calcium transients in these cells. To specifically ablate OTRs in CeL astrocytes, an rAAV vector allowing expression of Cre recombinase under the control of the astrocyte-specific glial fibrillary acidic protein (GFAP) promoter (rAAV-GFAPp-GFP-IRES-Cre) was injected into the CeL of an OTR conditional knockout (OTR-cKO) mouse line in which *loxP* sites flank the OTR coding sequence^25^ (Figure 2c). The injection of a control vector (rAAV-GFAPp-GFP) in OTR-cKO mice did not alter the OTR expression pattern, as 15.9% of astrocytes remained positive for OTR mRNA (Figure 2c), a proportion similar as what was found in rats CeL (Figure 1a-c). In comparison, CeL injection of the rAAV-GFAPp-IRES-Cre vector significantly reduced the number of OTR mRNA-expressing astrocytes to 1.2% (Figure 2c, S2e) but did not alter the OTR mRNA signals in neurons (Figure S2f), thus resulting in the CeL astrocyte-specific genetic deletion of OTRs. We then performed calcium imaging in CeA slices of mice using the calcium indicator Rhod-2 and identified astrocytes through GFP fluorescence. In those animals, TGOT failed to evoke calcium transients in CeL astrocytes, which was in start contrast to what we observed in control mice (Figure 2d).

Altogether, these findings suggest that OTR-expressing CeL astrocytes are fully equipped to respond in an autonomous fashion to the activity of oxytocinergic axonal projections arising from the hypothalamus in both rats and mice.

One interesting discrepancy is that up to 50% of CeL astrocytes responded to evoked release of OT or bath application of TGOT (Figure 1f-h, 2a) while OTR mRNA was detected in only 13 to 15% of CeL astrocytes (Figure 1a-c, S1a, 2a-c). Astrocyte calcium responses are known to spread through the astrocytes network via gap-junctions or paracrine communication through ATP release^20^. We first applied purinergic antagonists, PPADS, Suramin, and A438079 but failed to reveal any changes in the proportion or magnitude of CeL astrocytes responses to TGOT (Figure 2b, S2d). This suggests that purinergic transmission is not involved in the sequential activation of the CeL astrocytes network after OTR activation. However, carbenexolone (CBX), a widely-used blocker of gap junctions, significantly decreased the proportion of CeL astrocytes responding to OTR activation (16.5%) without impairing the calcium response dynamics in the remaining responsive CeL astrocytes (Figure 2b).

These findings suggest that the initial OTR-induced activation of CeL astrocytes spreads across the CeL astrocytic network through gap junctions.

### CeL Astrocytes are Sufficient to Recruit the CeL→CeM Neuronal Circuit

To address the potential modulatory role of CeL astrocytes on the CeL→CeM neuronal circuit, we first asked whether CeL astrocyte activity was sufficient to solely modulate CeA neuronal circuits (Figure 3). Using rAAVs to express the light-gated opsin C1V1(t/t) under the control of the GFAP promoter (Figure 3a), we specifically targeted CeL astrocytes (Figure S3a) and tested the effect of light-evoked calcium transients elicited in CeL astrocytes on the CeL→CeM neuronal circuit (Figure 3b).

**Figure 3.**
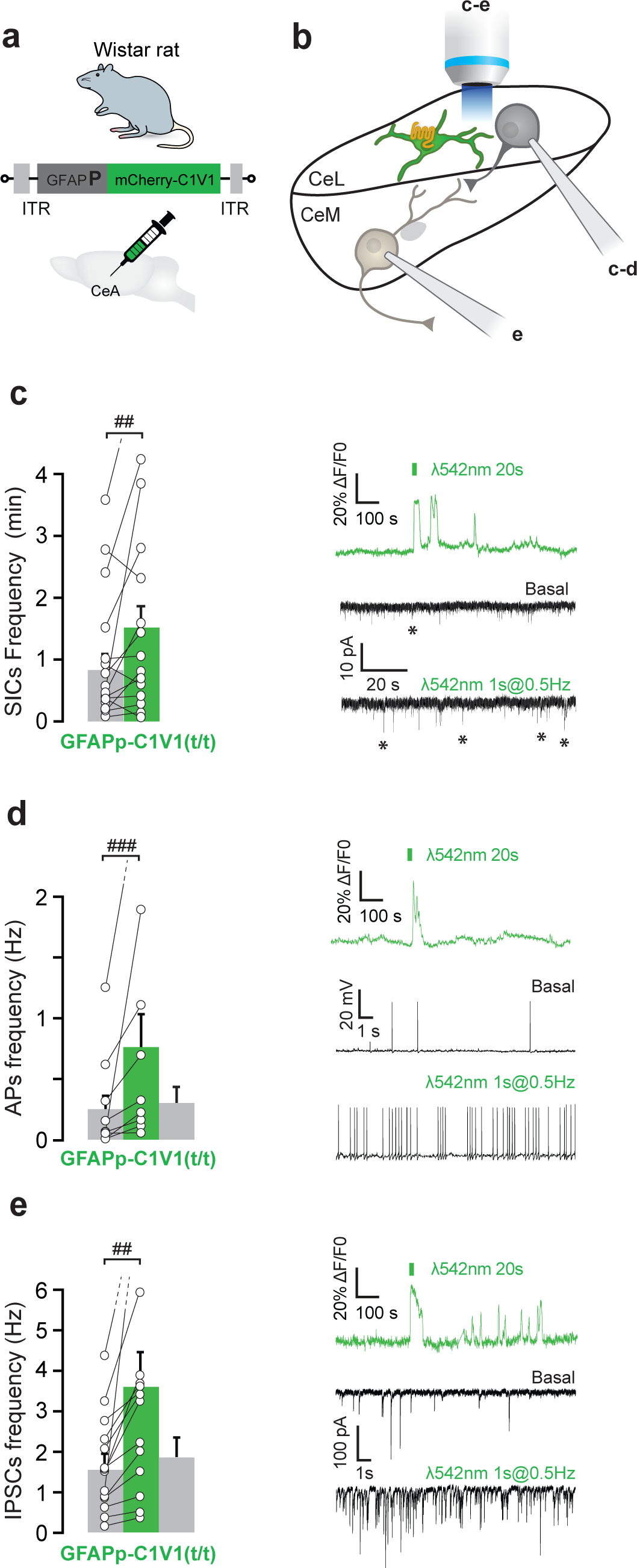
CeL Astrocytes Increased Activity is Sufficient to Recruit the CeL→CeM Neuronal Circuit. (**a**) Experimental rat model. (**b**) Scheme of the experimental setup, letters indicate in which figure subpart the technique was applied. (**c**-**e**) (Left) Effect of 3 min long light-evoked activation of C1V1 expressing astrocytes (1 s width, 0.5 Hz) on SICs and APs (**d**) frequencies recorded in CeL neurons (n = 19 and 10, respectively), and on (**e**) IPSCs frequencies recorded in CeM neurons (n = 19). (Top, right) Example traces of the effect of a 20 s, continuous, λ 542 nm light exposure on the ΔF/F0 of a C1V1 expressing astrocyte; (bottom, right) typical electrophysiological recordings traces. Patch-clamp data are expressed as averaged frequency plus SEM across cells before, during (and after) light effect; linked white circles indicate individual cell values. ^##^ *P* < 0.01, ^###^ *P* < 0.001 Wilcoxon signed rank test (Statistics in Table S3).

We hypothesized that a form of astrocytes-mediated neuromodulation could reveal itself through the presence of slow inward currents (SICs), a hallmark of neuronal NMDAR activation by astrocytes^26, 27^. SICs were indeed detected in CeL neurons (Figure 3c, S3b) and their frequency significantly increased after light-evoked calcium transients in CeL astrocytes (Figure 3c). To assess whether this increase in the SICs frequency is sufficient to increase excitability and ultimately firing in CeL neurons, we next performed *ex vivo* patch-clamp recordings of action potentials (APs) in CeL neurons upon CeL astrocytes stimulation (Figure 3b). We found that light-evoked calcium transients in CeL astrocytes led to an increase of APs frequencies in CeL neurons (Figure 3d). Crucially, this effect was prevented if we first loaded the Ca^2+^ chelator BAPTA into the CeL astrocytes syncytium^28^ (Figure S3c).

Given that the CeM is the main output of the CeA, we then conducted *ex vivo* patch-clamp recordings of inhibitory post-synaptic currents (IPSCs) in CeM projection neurons (Figure 3b), whose frequency is known to increase after CeL interneurons firing^3–5^. Indeed, we found that light-evoked calcium transients in CeL astrocytes led to an increase of IPSCs frequencies in CeM neurons (Figure 3e) which was also prevented by BAPTA infusion in the CeL astrocytes syncytium (Figure S3d). Altogether, these results indicate that CeL astrocytes activity is sufficient to modulate the CeL→CeM neuronal circuit.

### CeL Astrocytes are Necessary for the CeL→CeM Neuronal Circuit Modulation by OTR Activation

We then further examined whether CeL astrocytes are required to support the oxytocinergic modulation of the CeL→CeM neuronal circuit. We repeated the measurement of neuronal properties (Figure 3) and found that CeL neurons’ SICs and APs frequencies, as well as CeM neurons’ IPSCs frequency, were significantly increased after TGOT application in both rats and mice (Figure 4a1-a3, 4b1-b3; S4a).

Next, we loaded the Ca^2+^ chelator BAPTA into the rat CeL astrocytes syncytium (Figure 4a). BAPTA loading of CeL astrocytes significantly reduced the OTR-induced increase of SICs and APs frequencies in CeL neurons as well as the increase in IPSCs frequencies in CeM neurons (Figure 4a1-3). These results indicate the necessity of functional CeL astrocytes for the OTR-mediated recruitment of the CeL→CeM neuronal circuit.

**Figure 4.**
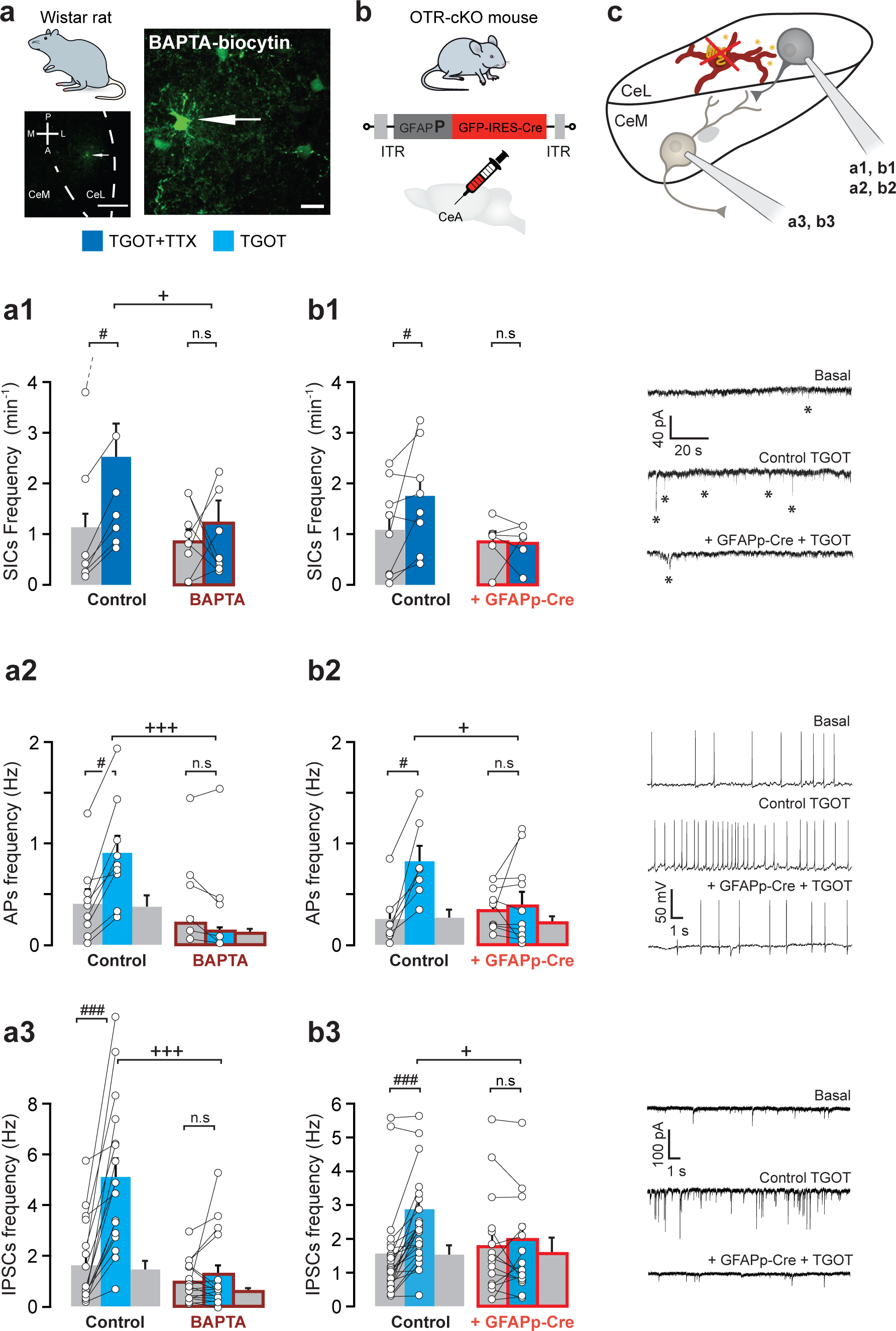
CeL Astrocytes are Necessary for the CeL→CeM Neuronal Circuit Modulation by OTR Activation. (**a**) Experimental rat model and experiments. (Bottom left) Localization of biocytin-filled astrocytes in the CeA rat brain horizontal slice. (Right) High magnification image. Arrow points to the primary filled cell. Scale bars 400 (left) and 50 µm (right). (**b**) Experimental mouse model and (**c**) scheme of the experimental setup. Letters indicate in which figure subpart the technique was applied: (**a1**-**3**) Consequences of BAPTA loading (40 mM, 45 min) into CeL astrocytes syncytium on TGOT-induced effect on SICs (**a1**, n = 7 and 8, dark blue) and APs (**a2**, n = 9 and 9, light blue) frequencies recorded in CeL neurons, and IPSCs frequencies (**a3,** n = 19 and 17) recorded in CeM neurons. (**b1**-**3**) (Left) Effects of TGOT (400 nM) on mice injected with GFAPp-GFP vector (control; left) or GFAPp-GFP-IRES-Cre vector (red borders; right) on SICs (**b1**, n = 8 and 5, dark blue) and APs (**b2**, n = 7 and 11, blue) frequencies recorded in CeL neurons, and IPSCs frequencies (**b3,** n = 27 and 16) recorded in CeM neurons. (Right) Typical electrophysiological recordings traces. Patch-clamp data are expressed as averaged frequency plus SEM across cells before, during (and after) drug effect; linked white circles indicate individual cell values. ^#^ *P* < 0.001, ^###^ *P* < 0.001 Wilcoxon signed rank test; ^+^ *P* < 0.05, ^+++^ *P* < 0.001 Mann-Whitney U test (Statistics in Table S4).

To further verify that the OTR-induced modulation of the CeL→CeM neuronal circuit depends on OTRs present on CeL astrocytes, we specifically ablated OTRs in CeL astrocytes of OTR-cKO mice (Figure 2c, S2e-f, 4b). In accordance with our previous results using pharmacological inactivation of astrocytes in rats, the specific deletion of OTR in mice CeL astrocytes significantly decreased the TGOT-induced increase of both SICs and APs frequencies in CeL neurons, as well as in IPSCs frequency in CeM neurons (Figure 4b1-3). These results highlight the necessity of OTR expression in CeL astrocytes for the OTR-mediated recruitment of the CeL→CeM neuronal circuit.

### CeL Astrocytes Recruit the CeL→CeM Neuronal Circuit through NMDAR modulation

We then sought to confirm that the activation of neuronal NMDAR, which is underlying the presence of SICs^26, 27^, is indeed a key component of CeL astrocyte-to-CeL neuron communication (Figure 5a-b). While the blockade of NMDAR with (2R)-amino-5-phosphonopentanoate (AP5, glutamate site) did not alter CeL astrocytic responses to TGOT (Figure S5a), AP5 prevented TGOT-induced modulation of SICs and APs frequencies in CeL neurons (Figure 5c-d). Moreover, OTR-induced CeL astrocyte-mediated SICs were linked to the activation of NR2B containing NMDARs, since ifenprodil prevented only the TGOT-induced SICs frequency increase (Figure 5c). We next tested whether repeated TGOT applications consistently modulate the CeL→CeM neuronal circuit via OTR-specific activation to assess the role of the NMDAR pathway in TGOT responsive CeM neurons (Figure S5b). We found that both AP5 and 5,7-dichlorokynurenic acid (DCKA, a blocker of the NMDAR glycine co-agonist site) significantly reduced the TGOT-evoked increase of IPSCs frequencies in CeM neurons (Figure S5c). On the contrary, the blockade of the α-amino-3-hydroxy-5-methyl-4-isoxazolepropionic acid receptor (AMPAR) had only a limited effect (Figure S5d). These results indicate that the oxytocin signal is directly conveyed from CeL astrocytes to CeL neurons via neuronal NMDAR activation.

**Figure 5.**
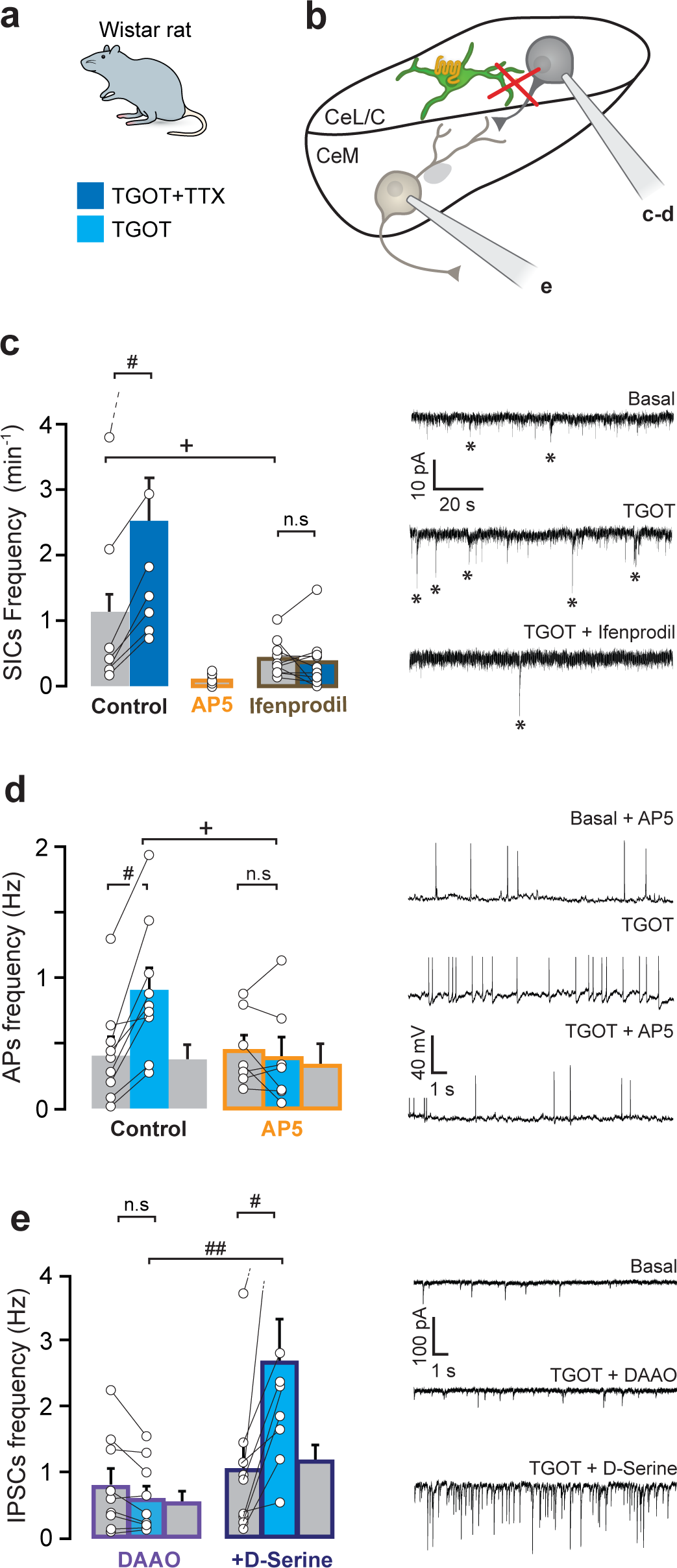
CeL Astrocytes Recruit the CeL→CeM Neuronal Circuit through NMDAR modulation. (**a**) Experimental rat model. (**b**) Scheme of the experimental setup. Letters indicate in which figure subpart the technique was applied. (**c**) Effect of TGOT (400nM; n = 7), TGOT+AP5 (50 µM; n = 10; orange borders) or TGOT+ifenprodil (3 µM; n = 10; brown borders) on SICs frequency recorded in CeL neurons. (**d**) Effect of TGOT (400 nM; n = 9) and TGOT+AP5 (50 µM; n = 7; orange borders) on APs frequencies recorded in CeL neurons. (**e**) Effect of DAAO (0.15 U/ml, incubation time > 1h30; purple borders), followed by incubation in D-Serine (20 min, 100 µM; dark purple borders) on TGOT-induced increase of IPSCs frequencies in CeM neurons (n = 9). Patch-clamp data are expressed as averaged frequency plus SEM across cells before, during (and after) drug effect; linked white circles indicate individual cell values. ^#^ *P* < 0.001, ^##^ *P* < 0.01 Wilcoxon signed rank test; ^+^ *P* < 0.05 Mann-Whitney U test (Statistics in Table S5).

We next sought to identify the molecular agent released by CeL astrocytes that gates the increase in CeL neuronal NMDAR activation. We used D-amino-acid oxidase (DAAO) to selectively catabolize the NMDAR co-agonist D-serine, a major astrocytic gliotransmitter^29–31^. TGOT had no effect after DAAO incubation, but subsequent incubation of the same cells in D-serine-containing ACSF partially restored the TGOT-mediated increased of IPSCs frequencies in CeM neurons (Figure 5e). Altogether, these data suggest that OTR-mediated activation of CeL astrocytes leads to the release of D-serine, allowing an increased activation of NMDAR in CeL GABAergic interneurons which, by increasing these neurons firing rate, leads in an increase of IPSCs frequencies downstream of the circuit, in CeM projection neurons.

### CeL Astrocytes are Sufficient to Modulate CeA Behavioral Correlates of Positive Emotional State and are Necessary for their OTR-mediated Modulation

Finally, we aimed to determine whether the CeL astrocyte-mediated modulation of the CeL→CeM neuronal circuit by OTR activation contributes to known CeA behavioral correlates. Since the CeA is a key structure in the regulation of pain-associated disorders^32^ and assigns emotionality to salient external stimuli^33^, we first tested the effect of TGOT on increased pain sensitivity and anxiety exhibited by neuropathic rats 4 weeks after a spared nerve injury surgery procedure^34, 35^ (SNI; Figure 6a). As neuropathy may cause synaptic changes in the amygdala^7, 36^, we conducted *ex vivo* patch-clamp recordings of IPSCs in CeM projection neurons 4 weeks after SNI or sham surgery. The TGOT-induced increase of IPSCs frequency was strictly similar in both conditions (Figure S6a4), indicating that the OTR-induced modulation of the CeL→CeM neuronal circuit is not altered after induction of SNI. Cannulae-guided micro-infusion of TGOT into the CeL slightly reduced mechanical hyperalgesia (Figure 6a1, S6a1) and clearly reduced anxiety levels in neuropathic animals (Figure 6a2, S6a2). We then performed similar experiments on SNI animals in which C1V1(t/t) was expressed under the control of a GFAP promoter in the CeL. Light-evoked stimulation of CeL astrocytes did not visibly affect mechanical pain thresholds of neuropathic animals, but was sufficient to reduce their anxiety (Figure 6a1-a2, S6a1-a2). Since the amygdala is critically involved in the regulation of emotions^37^, we performed a conditioned place preference test (CPP) to assess if the OTR-induced modulation of the CeL→CeM pathway is linked to the emotional component of pain. Both cannulae-guided micro-infusion of TGOT in CeL and light-evoked activation of C1V1(t/t)-expressing CeL astrocytes induced a strong place preference in neuropathic and sham-operated rats (Figure 6a3, S6a3).

**Figure 6.**
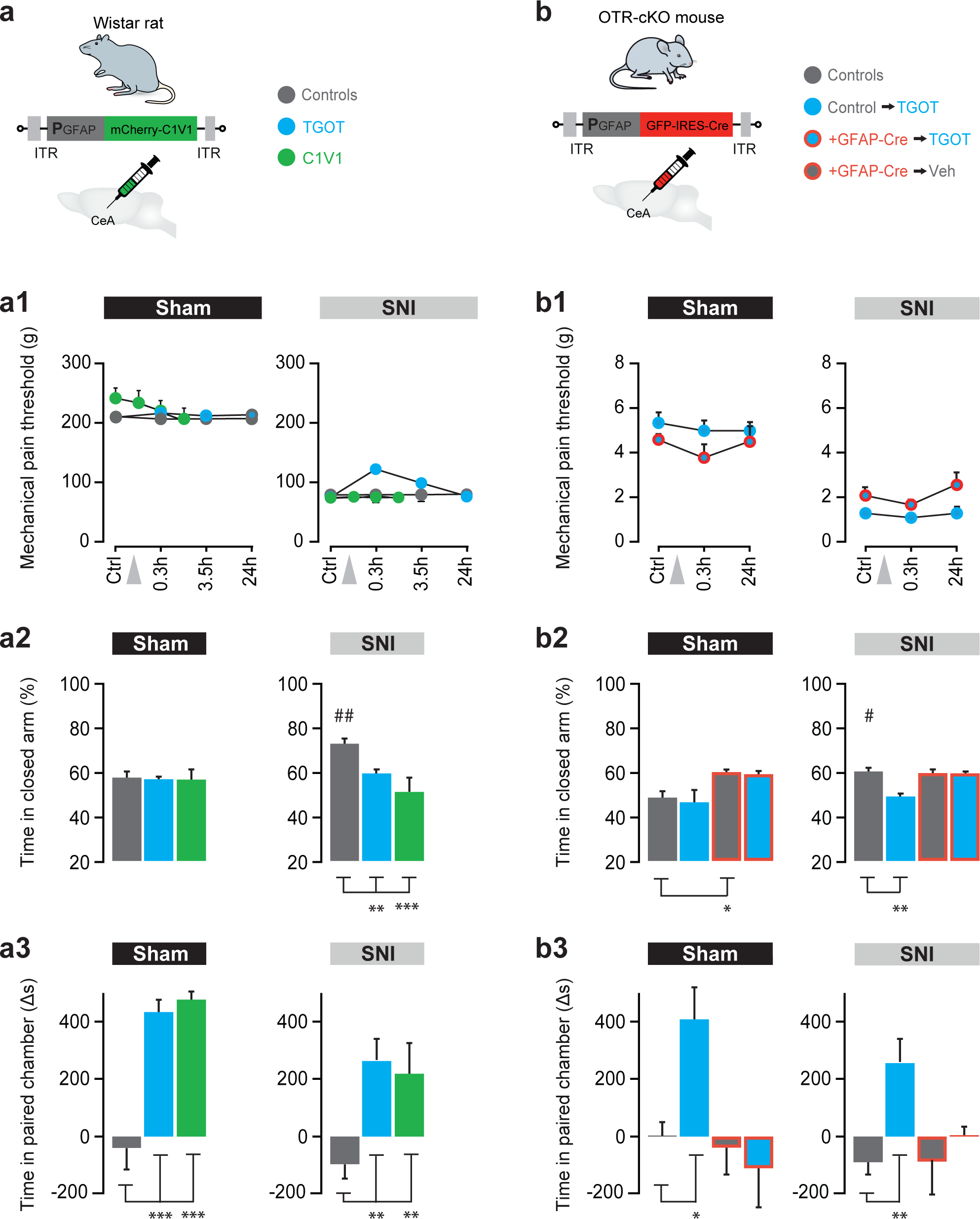
CeL Astrocytes are Sufficient to Modulate CeA Behavioral Correlates of Comfort and Necessary to their OTR-mediated Modulation. (**a**) Experimental rat model. TGOT or vehicle were injected 20 min before behavioral tests. rAAV-GFAPp-C1V1(t/t)-mCherry was injected 2 weeks before tests, astrocytes were light-stimulated for 3 minutes with 1 s width pulse at 0,5 Hz immediately prior measurements. (**b**) Experimental mice model. TGOT or its vehicle were administered in OTR-cKO mice previously injected within the CeL either with rAAV-GFAPp-GFP (controls) or rAAV-GFAPp-GFP-IRES-Cre 4 weeks before tests. (**a1**, **b1**) 4 weeks after the SNI surgery, mechanical pain threshold was assessed on the neuropathic paw before (Ctrl) and after either drugs injections or C1V1 light-driven activation of CeL astrocytes (gray arrow) for sham and SNI animals. (**a2**, **b2**) Anxiety levels were assessed through measurements of the time spent in the closed arms of the elevated plus maze after drugs injections or C1V1 light-driven activation of CeL astrocytes for sham and SNI animals. (**a3**, **b3**) Conditioned place preference (CPP) was assessed through measurements of the Δ time spent in the paired chamber before and after pairing. Pairing was realized through drugs injections or C1V1 light-evoked activation of CeL astrocytes for SNI and sham animals. Data are expressed as averages across rats plus SEM. n = 4-18 per group (details in Table S6). ^#^ *P* < 0.001, ^##^ *P* < 0.01 Wilcoxon signed rank test; * *P* < 0.05, ** *P* < 0.01, *** *P* < 0.001; ANOVA or mixed-design ANOVA followed by posthoc Bonferroni test (Statistics in Table S6).

To confirm that the OTR-induced modulation of the observed behaviors strictly depends of OTR present on CeL astrocytes, we specifically ablated OTRs in mice CeL astrocytes (Figure 2c, S2e-f, 6b) and tested the effect of TGOT on increased pain sensitivity and anxiety exhibited by neuropathic mice 4 weeks after a spared nerve injury surgery procedure^38^ (SNI; Figure 6b). Cannula-guided micro-infusion of TGOT in CeL did not change mechanical hyperalgesia (Figure 6b1, S6b1) but reduced anxiety levels in control neuropathic animals (Figure 6b2, S6b2), an effect that was prevented by the CeL astrocyte specific deletion of OTRs (Figure 6b2). As previously shown in rats (Figure 6a3), we tested mice for CPP and found a strong place preference effect of micro-infusion of TGOT in CeL in both neuropathic and sham-operated control mice, which was absent after deletion of OTR in CeL astrocytes (Figure 6b3, S6b3).

Altogether, these findings point towards the OTR-mediated signaling in CeL astrocytes as a counterbalancing mechanism supporting the positive emotional valence under both chronic pain and healthy states. It indicates that the OTR-astrocytes mediated modulation of the CeL→CeM neuronal circuit is not only involved in the regulation of hyperalgesia or anxiety but also represents a more general cellular substrate underlying the promotion of a positive emotional state (Figure 7).

**Figure 7.**
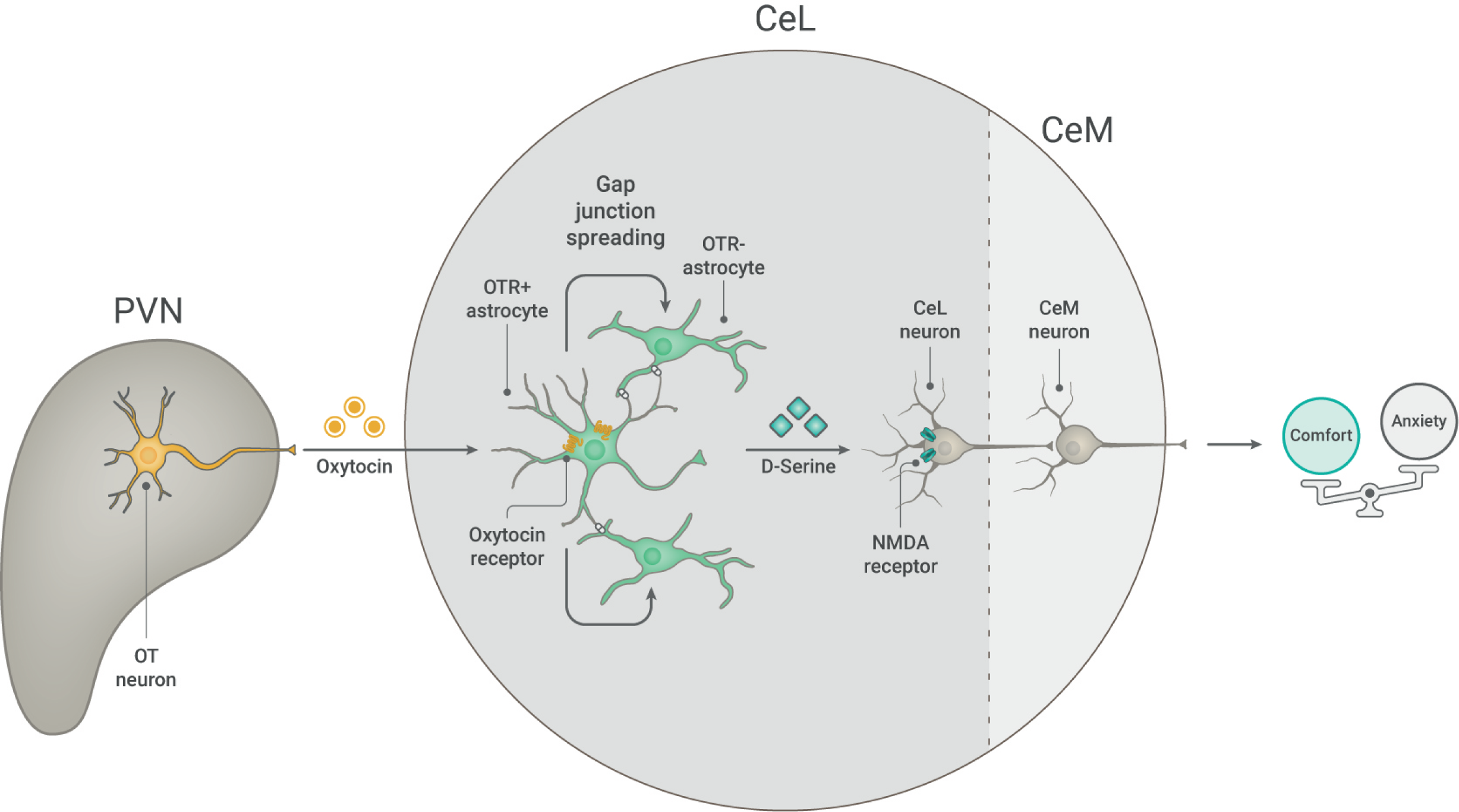
Oxytocin-dependent Interactions in the CeA. Based on our findings we hypothesize that oxytocin released from axons of PVN oxytocinergic neurons within the CeL activates the oxytocin receptor (OTR)-expressing astrocytes and their neighboring astrocytes inter-connected via gap junctions. Subsequently, the CeL astrocytes release D-serine which gate the activation of CeL interneurons, in turn inhibiting CeM output projection neurons, resulting in anxiolysis and the promotion of a positive emotional state.

## DISCUSSION

In addition to their classical functions related to structural and metabolic support, modulation of synaptic transmission and transmitter uptake and release, astrocytes have emerged as key players in intricate and diverse glia-neuronal network interactions^19^. However, the functional relevance of astro-neuronal interactions in the modulation of complex social and emotional behaviors remains elusive. Following the initial experiments on OTRs in cultured astrocytes^15^, we sought to investigate the role of these cells in OTR signaling in the CeA, a structure critically involved in the processing of emotionally-relevant information. We showed that CeL astrocytes play a critical role in the oxytocin-induced modulation of the CeL→CeM neuronal circuit in two rodent species, rats and mice. Here we report that a subpopulation of CeL astrocytes express functional OTR and convey the oxytocin signal among other (OTR-negative) astrocytes of the CeL through gap junctional coupling. This recruitment of the CeL astrocytes syncytium leads to an NMDAR-mediated tuning of CeA neuronal circuits. At the behavioral level, this oxytocin-mediated modulation of the astro-neuronal network of the central amygdala might promote a positive emotional state.

The amygdala is a key region for emotional processing^37^ with a clear involvement in pain regulation and particularly persistent pain, which impacts ∼8% of the global population^7^. On the other hand, oxytocin is a neuropeptide well known to act on various behaviors throughout the cycle of life, with major involvement in a broad variety of physiological functions, all directed toward well-being, emotional balance and ultimately species survival^1, 39^. In this regard, it is interesting to observe that our results suggest that the oxytocin-induced modulation of CeA networks not only attenuates hypersensitivity symptoms related to nociception in itself, but more generally improves the positive emotional valence of the animals even under pain-free conditions. Pain management is a clinically difficult topic, and most of the newly developed analgesic drugs have failed or not met expectations and patients’ needs. Therefore, it might be particularly relevant to consider new therapeutic strategies aiming at alleviating the pain spectrum globally, for example by targeting the oxytocin system, to improve the overall patient well-being.

The oxytocinergic regulation of neural circuits and oxytocin’s physiological functions is presently under intense scrutiny^6, 40–46^. Yet, to the best of our knowledge, these studies were solely based on the hypothesis that oxytocin signaling occurs predominantly in neurons, despite a sparse number of independent studies demonstrating expression of OTR in glia, notably astrocytes^13, 15, 16, 47, 48^. Here, we propose a novel mechanism implicating astrocytes as key contributors of oxytocin action. The induction of long-lasting calcium oscillations in astrocytes following OTR activation leads to a sustained modulation of the CeL→CeM neuronal circuit, affecting known CeA functions such as emotional valence regulation. It may seem counter-intuitive that without a pristine astroglial function, CeA neurons appear unaffected by OTR activation, despite their clear transcription of OTR mRNA. We hypothesize that OTR-induced activation of CeL astrocytes gates CeL neurons responses to oxytocin, and probably other synaptic inputs, through sustained (co-) activation of NMDARs. Such mechanism of astrocytes to neuron communication through activation of extra-synaptic NMDARs has also been proposed by other studies to favor a synchronous increase in excitability across an ensemble of neurons^49, 50^. In particular, activation of Gq or Gi/o G-protein coupled receptors (GPCRs) specifically in astrocytes enhances neuronal excitability and ultimately firing^51^, a result which could parallel the activation of the same signaling pathways here by specific OTR activation, and GPCRs are known to be coupled to either of Gq or Gi/o proteins^52^. This proposed mechanism would allow for a synchronized and long-lasting switch in the gain of the CeA neuronal circuits, thereby amplifying the effect of oxytocin on CeA outputs in both the spatial and temporal domains. In light of the predominantly non-synaptic OT release from axons *en passant*, which could lead to CeL-restricted micro-volume transmission of the neuropeptide^53, 54^, it seems plausible that astrocytes relay and amplify oxytocin signaling to neurons. This would provide control over the oxytocin effect on neuronal circuits, as has already been proposed for other neuromodulators and neurotransmitters^55, 56^. The different dynamics and connectivity of astro-neuronal networks provide new insights in the regulation of brain functions and ultimately animal behavior^57^, with a complexity yet to be unraveled^20, 22, 58^. From a larger perspective, our results further confirm that astrocytes are functionally diverse and can regulate central nervous system functions in a network-specific manner^19, 59, 60^. Therefore, the study of astrocyte-neuron interactions is essential for a better comprehension of both physiological and pathological mechanisms affecting brain function^20, 60, 61^.

Given that oxytocin modulates emotions and behaviors through a variety of actions on the peripheral and central nervous system^2, 62–64^, our study adds a novel mechanism of oxytocin action: the modulation of astrocyte-neuron interactions. This mechanism underpins the oxytocinergic promotion of a positive emotional state, opening up new perspectives for studying the role of astroglial-neuronal networks in the modulation of emotional experiences^57, 65^, notably by neuropeptidergic systems^66^. On a broader scope, our present results demonstrate that the astroglia-mediated effects of oxytocin provide a new basis for the exploration of pathophysiologic mechanisms and enlighten a new vision of interplay between neuropeptides and glia that bears a general value for a better understanding of functional brain organization. This opens up new possibilities of research about the neuronal/glial substrates underlying positive emotional states, which is highlighted in our working model (Figure 7). The development of prospective treatments for somatic and mental diseases in humans may indeed be helped by targeting the oxytocin or other neuromodulators systems, and their joint action on multiple CNS cell types.

## ACKNOWLEDGEMENTS

This work was supported by the IASP Early Career Research grant 2012, FP7 Career Integration grant 334455, Initiative of Excellence (IDEX) Attractiveness grant 2013, IDEX Interdisciplinary grant 2015, University of Strasbourg Institute for Advanced Study (USIAS) fellowship 2014-15, Foundation Fyssen research grant 2015, NARSAD Young Investigator Grant 24821, ANR JCJC grant (to AC); ANR-DFG grant GR 3619/701 (to AC and VG); the Schaller Research Foundation, DFG grant GR 3619/13-1, DFG within the Collaborative research Center SFB 1134, (to VG) and SFB 1158 (to CP and VG); SNSF-DFG grant GR 3619/8-1 (to RS and VG), Fritz Thyssen foundation (to VG); Alexander von Humboldt Foundation (to DH); Fyssen foundation and PROCOP grant and SFB1158 seed grant for young scientists (to AL); Research Foundation - Flanders, fellowship (12V7519N) (to JW); Russian science foundation RSF (17-75-10061) and the Program of Competitive Growth of Kazan University (to AR); the intramural research program of the NIMH (ZIAMH002498) (to WSY); National Institutes of Health grants R01NS094640 and R01HL090948 (to JES). The authors thank Vincent Lelièvre for *in situ* hybridization advices; Romain Goutagny for *in vivo* optogenetics assistance; Fulvio Magara for anxiety behavior advices; Barbara Kurpiers and the Interdisciplinary Neurobehavioral Core Facility of Heidelberg University for experiments performed there; Sophie Reibel, Dominique Ciocca and the Chronobiotron UMS 3415 for all animal care; Thomas Splettstoesser (www.scistyle.com) for its initial help with the preparation of figures.

## AUTHOR CONTRIBUTIONS

Conceptualization, AC; Methodology, AC, BBo, CML, CP, DK, FA, ID, JW, JYC, PD, PP, RS, VG, WSY, YG; Analysis, AC, BBe, BBJ, CML, DK, FA, HSKB, JW, SG; *In situ* hybridization, DH, FA, HSKB, HW, JS, ME; Immunohistochemistry, AL, DH, FA, JW, ME, MdSG; *Ex vivo* patch-clamp electrophysiology, AB, AC, JW, SG, DK, IW, BBe, MA; *Ex vivo* calcium imaging, AB, CML, DK, JW; Astrocytes characterization, AR, DK, IW, ME, SG; Behavior, AC, BBJ, DK, JW; Mice line validation, WSY; Viral vectors validation, MdSG, ME, VG; Spared nerve injuries, PI, MP; Writing, AC, BBo, JES, JYC, JW, RS, VG; Funding acquisition AC, VG; Supervision, AC; Project administration, AC.

## MATERIALS AND METHODS

### KEY RESOURCES TABLE

All the reagents used are listed in the Supplementary Table 7

### CONTACT FOR REAGENTS AND RESOURCE SHARING

Further information and requests for resources and reagents should be directed to and will be fulfilled by the Lead Contact, Alexandre Charlet. The requests for viral vectors should be addressed to Valery Grinevich.

### EXPERIMENTAL MODEL AND SUBJECT DETAILS

#### Animals

Animals were housed under standard conditions with food and water available *ad libitum* and maintained on a 12-hour light/dark cycle and all experiments were conducted in accordance with EU rules and approbation from French Ministry of Research (01597.05). For *ex vivo* and *in vivo* experiments, Wistar rats or C5BL/6 mice were used. *Ex vivo* experiments used animals between 18 and 25 days old, except in experiments where rAAVs were injected, in which case animals were between 2 and 6 months old at the time of sacrifice. *In vivo* experiments used 2-month-old animals at the time of the first surgery.

#### Cloning and Production of rAAV Vectors

The generation of rAAVs allowing for the specific expression of the protein of interest in OT-cells is described in our previous work ^5^. Briefly, the conserved promoter region of 2.6 kb was chosen using the software BLAT from UCSC (http://genome.ucsc.edu/cgi-bin/hgBlat), was amplified from BAC clone RP24-388N9 (RPCI-24 Mouse, BACPAC Resources, CHORI, California, USA) and was subcloned into a rAAV2 backbone carrying an Ampicillin-resistance.

To construct the OTp-C1V1(t/t)-TS-mCherry AAV vector we used previously cloned OTp-DIO-GFP-WRE plasmid^2^ equipped with the characterized 2.6 kb OT promoter^5^. In this plasmid the DIO-GFP sequence was replaced by C1V1(t/t)-TS-mCherry from the rAAV CaMKIIa-C1V1(t/t)-TS-mCherry (Addgene, plasmid #35500).

To generate GfaP-C1V1(t/t)-TS-mCherry AAV vector, we replaced the CamKIIa promoter from the rAAV CaMKIIa-C1V1(t/t)-TS-mCherry by the Gfa promoter from the pZac2.1 gfaABC1D-tdTomato (Addgene, plasmid: 44332). The cell type specificity of the rAAV carrying the Gfa promoter was recently confirmed^67^. In analogy, the generation of the GFAP-GFP-IRES-Cre vector was achieved using pZac2.1 gfaABC1D-tdTomato (Addgene, plasmid: 44332). First, the promoter was cloned into a rAAV2 backbone and sticky ends were blunted with EcoR1 and Basrg1. Next, pAAV-CamKIIa-C1V1(t/t)-TS-mCherry was blunted using BamHI and BsrGI. Finally, the pBS-ires cre construct was used and IRES-Cre was inserted into the GFAP-driven vector resulting in the GFAP-GFP-IRES-Cre construct.

Production of chimeric virions (recombinant Adeno-associated virus 1/2; rAAV 1/2) was described in^5^. Briefly, human embryonic kidney cells 293 (HEK293; Agilent #240073) were calcium phosphate-transfected with the recombinant AAV2 plasmid and a 3-helper system. rAAV genomic titers were determined with QuickTiter AAV Quantitation Kit (Cell Biolabs, Inc., San Diego, California, USA) and are ∼10^^13^ genomic copies per ml for all rAAV vectors used in this study.

## METHOD DETAILS

### Surgeries

#### Neuropathic Pain Model: Spared Nerve Injury (SNI) Procedure

Animals were randomly separated in two groups to undergo either posterior left hindpaw SNI or sham procedure, with right hindpaw untouched. Animals were anaesthetized using isoflurane at 1.5–2.5%. Incision was made at mid-thigh level using the femur as a landmark and a section was made through the biceps femoris. The three peripheral branches (sural, common peroneal and tibial nerves) of the sciatic nerve were exposed. Both tibial and common peroneal nerves were ligated using a 5.0 silk suture and transected. The sural nerve was carefully preserved by avoiding any nerve stretch or nerve contact^68^. For animals undergoing sham surgery, same procedure was performed but nerves remained untouched. Animals were routinely observed daily for 7 days after surgery and daily tested by the experimenter (Figure S4). Besides observing weight, social and individual behavior, the operated hindpaw was examined for signs of injury or autotomy. In case of autotomy or suffering, the animal was euthanized in respect of the ethical recommendations of the EU. No analgesia was provided after the surgery in order to avoid interference with chronic pain mechanisms and this is in accordance with our veterinary authorization. Suffering was minimized by careful handling and increased bedding.

#### Stereotaxic Surgery: Injections of rAAV Vectors

Stereotaxic surgery was performed under deep ketamine-xylazine anesthesia, using the KOPF (model 955) stereotaxic system. For specific control of rats CeA astroglial cells, 200 nl of rAAV serotype 1/2 (GFAPp-C1V1TSmCherry, Cloned from plasmids #35500 and 44332, Addgene) were injected bilaterally at the coordinates corresponding to CeL: rostro-caudal: -2.7mm, medio-lateral: 4.2mm, dorso-ventral: -8.0mm (From Paxinos and Watson Atlas). For specific control of

OT neurons, 200 nl of rAAV serotype 1/2 (OTp-C1V1TSmCherry or OTp-ChR2mCherry) were injected bilaterally at the coordinates corresponding to each hypothalamic OT nuclei. PVN: rostro-caudal: -1.8mm; medio-lateral: +/-0.4mm; dorso-ventral: -8.0mm; SON: rostro-caudal: -1.4mm; medio-lateral: +/-1.6mm; dorso-ventral: -9.0mm; AN: rostro-caudal: -2mm; medio-lateral: +/- 1.2mm; dorso-ventral: -8.5mm (From Paxinos and Watson Atlas). For specific deletion of OTR in mice CeL astrocytes, 280 nl of rAAV serotype 1/2 (GFAPp-GFP-IRES-Cre) were injected bilaterally at the coordinates corresponding to CeL: rostro-caudal: -2.7mm, medio-lateral: 4.2mm, dorso-ventral: -8.0mm (From Paxinos and Watson Atlas) in OTR-cKO mice.

### Stereotaxic Surgery: intra-CeL Cannulae

#### Cannulae Implantation

Animals were bilaterally implanted with guide cannulae for direct intra-CeL infusions. As guide cannulae we used C313G/Spc guide metallic cannulae (Plastics one, VA, USA) cut 5.8 mm below the pedestal. For this purpose, animals were deeply anesthetized with 4% isoflurane and their heads were fixed in a stereotaxic frame. The skull was exposed and two holes were drilled according to coordinates that were adapted from brain atlas (rat, 2.3 mm rostro-caudal; 4 mm lateral; 7.5 mm dorso-ventral relative to bregma; mice, 1.4 mm rostro-caudal; 2.6 mm lateral; 4.3 mm dorso-ventral relative to bregma) by comparing the typical bregma-lambda distance with the one measured in the experimental animal. Two screws were fixed to the caudal part of the skull in order to have an anchor point for the dental cement. Acrylic dental cement was finally used to fix the cannulae and the skin was sutured. In case of long lasting experiments (neuropathy-induced anxiety) with a cannula implantation at distance of the behavioral assay (> 4 weeks), cannulae were sometimes lost or cloaked, and concerned animals therefore excluded from testing.

#### Drugs Infusions

We used bilateral injections of 0.5 μl containing either vehicle (NaCl 0.9%) or oxytocin agonist TGOT (1 μM) dissolved in NaCl 0.9%. For this procedure two injectors (cut to fit 5.8 mm guide cannulae protruding 2 to 2.5 mm beyond the lower end of the cannula in older animals and 1.8 mm in 3-4 week old rats) were bilaterally lowered into the guide cannula, connected via polyten tubing to two Hamilton syringes that were placed in an infusion pump and 0.5 μl of liquid was injected in each hemisphere over a 2-minute-period. After the injection procedure, the injectors were kept in place for an additional minute in order to allow a complete diffusion of liquid throughout the tissue. Rats were subsequently left in the home cage for 15 minutes to recover from the stress of the injection and then handled for mechanical pain threshold or anxiety assessment. Animals that received TGOT injections for the first experiment (mechanical sensitivity assessment) were switched to the vehicle injected groups for the elevated plus maze experiment.

### Stereotaxic Surgery: intra-CeL Optical Fiber

#### Optical Fiber Implantation

Sham and rAAVs injected animals both underwent a single surgical procedure in which after vector injection or no injection for sham, optical fibers designed to target the CeL were implanted and firmly maintained on the skull using dental cement. See “***cannulae implantation*”** for the surgical procedure. Implantable optical fibers were homemade using optical fiber cut at appropriate length (FT200EMT, Thorlabs, NJ, USA) inserted and glued using epoxy based glue in ferrules (CFLC230-10, Thorlabs, NJ, USA).

### Horizontal and Coronal Slices

#### Slices Preparations

In all cases, animals were anaesthetized using ketamine (Imalgene 90 mg/kg) and xylazine (Rompun, 10 mg / kg) administered intra-peritoneally. Transcardial perfusion was then performed using one of the following artificial cerebro-spinal fluids (ACSFs) dissection solutions. For animals between 18 and 25 days old, an ice-cold sucrose based dissection ACSF was used containing (in mM): Sucrose (170), KCl (2.5), NaH2PO4 (1.25), NaHCO3 (15), MgSO4 (10), CaCl2 (0.5), HEPES (20), D-Glucose (20), L-ascorbic acid (5), Thiourea (2), Sodium pyruvate (3), N-acetyl-L-cysteine (5), Kynurenic acid (2). For animals between 2 and 6 months old, an ice-cold NMDG based ACSF was used containing (in mM): NMDG (93), KCl (2.5), NaH2PO4 (1.25), NaHCO3 (30), MgSO4 (10), CaCl2 (0.5), HEPES (20), D-Glucose (25), L-ascorbic acid (5), Thiourea (2), Sodium pyruvate (3), N-acetyl-L-cysteine (10), Kynurenic acid (2). In both cases, pH was adjusted to 7.4 using either NaOH or HCl, this after bubbling in 95% O2-5% CO2 gas, bubbling which was maintained throughout the duration of use of the various ACSFs. Those ACSFs formulae were based on the work of^69^. Following decapitation, brain was swiftly removed in the same ice-cold dissection ACSFs as for transcardial perfusion, and 350 µm thick horizontal slices containing the CeA obtained using a Leica VT1000s vibratome. For experiments in Figure S1E, coronal slices of the same thickness containing the PVN were used. Upon slicing, brain slices were hemisected and placed, for 1 hour minimum before any experiments were conducted, in a room tempered holding chamber, containing normal ACSFs. For 2 to 6 month old animals, slices were first let for 10 minutes in 35°C NMDG ACSF before placing them in the holding chamber at room temperature. Normal ACSF, also used during all *ex vivo* experiments, is composed of (in mM): NaCl (124), KCl (2.5), NaH2PO4 (1.25), NaHCO3 (26), MgSO4 (2), CaCl2 (2), D-Glucose (15), adjusted for pH values of 7.4 with HCL or NaOH and continuously bubbled in 95% O2-5% CO2 gas. All ACSFs were checked for osmolality and kept for values between 305-310 mOsm/L. In electrophysiology or calcium imaging experiments, slices were transferred from the holding chamber to an immersion recording chamber and superfused at a rate of 2 ml/min with normal ACSFs unless indicated otherwise.

#### Drug Application

OTR agonists were bath applied through a 20s long pumping of agonist solution, corresponding to several times the volume of the recording chamber. Other drugs (antagonists, TTX, channel blockers etc.) were applied for at least 20 minutes in the bath before performing any experiments. BAPTA (or BAPTA-free solution for controls) loading of CeL astrocytes was realized following^70^ protocol. Two distant CeL astrocytes per slice (label with SR101, 1 µM) were patched in whole cell configuration and voltage steps were applied (2 Hz, Δ40 mV) to help loading the BAPTA contained in the patch pipette (in mM): MgCl2 (1), NaCl (8), ATP (2) GTP (0.4) HEPES (10), BAPTA (40) and osmolality checked to be between 275-285 mOsm/l. The whole cell configuration was maintained during 45 min to allow BAPTA diffusion into the astrocytes syncytium.

### Calcium Imaging and Identification of Astrocytes

To identify astrocytes, SR101 (1 μM) was added to ACSF in a culture well and slices were incubated for 20 minutes at 35°C. The specificity of SR101 labelling to astrocytes of the CeL was verified through patch-clamp experiments, the results of which can be found in Figure S2a-c. After SR101 loading, the synthetic calcium indicator OGB1 or Rhod-2 was bulk loaded following an adapted version of the method described previously^69^ reaching final concentrations of either 0.0025 % (∼20 μM) for OGB1 or 0.0025 % (∼20 μM) for Rhod-2, 0.002% Cremophor EL, 0.01 % Pluronic F-127 and 0.5% DMSO in ACSF, and incubated for 45 to 60 minutes at 38°C. Upon incubation time, slices were washed in ACSF for at least an hour before any recording was performed. Astrocytes recorded for this study were then those co-labeled, in rats for SR101 and OGB1 and in mice for GFP and Rhod2. The spinning disk confocal microscope used to perform astrocyte calcium imaging was composed of a Zeiss Axio examiner microscope with a 40x water immersion objective (numerical aperture of 1.0), mounted with a X-Light Confocal unit – CRESTOPT spinning disk. Images were acquired at 2Hz with either a Rolera em-c² emCCD or an optiMOS sCMOS camera (Qimaging, BC, Canada). Cells within a confocal plane were illuminated for 100 to 150 ms for each wavelength (SR101 and Rhod-2: 575 nm, OGB1 and GFP: 475 nm) using a Spectra 7 LUMENCOR. The different hardware elements were synchronized through the MetaFluor software (Molecular Devices, LLC, Ca, USA) which was also used for online and offline quantitative fluorescence analysis. Astrocytic calcium levels were measured in hand drawn ROIs comprising the cell body plus, when visible, proximal processes, without distinction. In all recordings, the Fiji rolling ball algorithm was used to increase signal/noise ratio.. Absolute [Ca^2+^]i variation were estimated as changes in fluorescence signals over the baseline (ΔF/F). Baseline was established for each ROI as the average fluorescence over all pictures. To take into account micro-movements of the specimen on long duration recordings, the ΔF/F values were also calculated for SR101 / GFP and subsequently subtracted to the ones of OGB1 / Rhod2, except in the case of Figure 1f-h, where astrocytes were identified through SR101 fluorescence after the recordings, to avoid unwanted stimulation of the C1V1(t/t) opsin. On this last case, recordings in which movements / drifts were visible were discarded. Bleaching was further corrected by fitting a linear regression on the overall ΔF/F trace for each ROI, which values were then subtracted to the ΔF/F. Upon extraction of data, calculations and corrections of ΔF/F for each astrocyte, the area under the curve before and after OTR agonists’ application (or light pulse for optogenetics experiments) were calculated and proportionally corrected relative to the different baseline and after stimulation recording durations. An astrocyte was considered as being responsive if the relative ratio of AUCs after drug/light application over baseline was higher or equal to a 20% increase. The relative AUCs ratios values of responding astrocytes were used for quantitative analysis and called “relative AUC increase”, and in corresponding histograms the coordinate indicated as a 0 corresponds to a ratio of 1 (meaning neither increase nor decrease of the AUC). The maximal peak reached after drug/light application was also measured in responsive cells and used in quantitative analysis shown in various figures. Data were averaged across all responding astrocytes per slice, which were used as the statistical unit. For the response probability, calcium events were detected as an increases of fluorescence > 5% with a rising slope > 0,025 and a minimum inter-peak distance of 10 seconds. Then, the response probability was calculated as the number of astrocyte with a least 1 calcium event per time bin (30s) divide per the number of astrocyte in each recording. Image J software was also used on SR101 / OGB1 pictures to produce illustrative pictures. Finally, all calcium imaging experiments were conducted at controlled room temperature of 26°C.

### Electrophysiology

Whole cell patch-clamp recordings of CeL neurons, CeL astrocytes and CeM neurons were visually guided by infrared oblique light visualization of neurons and completed by SR101 fluorescence observation for astrocytes. Patch-clamp recordings were obtained with an Axon MultiClamp 700B amplifier coupled to a Digidata 1440A Digitizer (Molecular Devices, CA, USA). Borosilicate glass electrodes (R = 3.5 -7 MΩ) with inner filament (OD 1.5 mm, ID 0.86 mm; Sutter Instrument, CA USA) were pulled using a horizontal flaming/brown micropipette puller (P97; Sutter Instrument, CA, USA). Recordings were filtered at 2 kHz, digitized at 40 kHz and stored with the pClamp 10 software suite (Molecular Devices; CA, USA). Analysis of patch-clamp data were performed using Clampfit 10.7 (Molecular Devices; CA, USA) and Mini analysis 6 software (Synaptosoft, NJ, USA) in a semi-automated fashion (automatic detection of events with chosen parameters followed by a visual validation).

#### Whole-Cell Recording of CeL Neurons and Astrocytes

Recording pipettes were filled with an intracellular solution containing (in mM): KMeSO4 (125), CaCL2 (2), EGTA (1), HEPES (10), ATPNa2 (2), GTPNa (0.3). The pH was adjusted to 7.3 with KOH and osmolality checked to be between 290-295 mOsm/l, adjusted with sucrose if needed. All neurons were hold at a membrane potential of -65 mV, astrocytes -80 mV. Series capacitances and resistances were compensated electronically throughout the experiments using the main amplifier. For SICs measurements in CeL neurons (Figure 3, 4, S3, S4), whole cell recordings were conducted in a Mg^2+^ free ACSF, also containing biccuculin (10 µM) and TTX (1 µM) as in^27^. Currents were categorized as SIC when their rise time (10% to 90%) and decay tau were strictly superior to 8 and 30 ms, respectively. Average SICs frequencies per cell was calculated over 10 minutes periods during basal condition and 10 minutes periods after the bath application of TGOT. CeL neurons were classified as TGOT-responsive when the average SIC frequency was increased by at least 30% after TGOT application when compared to its baseline.

#### Whole-cell Recording of CeM Neurons

Pipettes were filled with an intracellular solution containing (in mM): KCl (150), HEPES (10), MgCl2 (4), CaCl2 (0.1), BAPTA (0.1), ATP Na salt (2), GTP Na salt (0.3). pH was adjusted to 7.3 with KOH and osmolality checked to be between 290-295 mOsm/L, adjusted with sucrose if needed. All cells were hold at a membrane potential of -70 mV. Series capacitances and resistances were compensated electronically throughout the experiments using the main amplifier. Average events frequencies per cell were calculated on 20s windows, chosen for TGOT or light stimulation during maximal effect, as determined by the visually identified maximal slope of the cumulative plot of the number of events. CeM neurons were classified as TGOT-responsive when the average IPSCs frequency was increased by at least 20% during at least 10s and up to 500s after TGOT application when compared to baseline average frequency. Baseline and recovery frequencies were measured respectively at the beginning and end of each recording. All patch-clamp experiments were conducted at room temperature.

### Immunohistochemistry and *in situ* Hybridization

#### In situ Hybridization for OTR mRNA in Rat CeL

The probe for OTR mRNA was *in vitro* transcripted from a 902-bp fragment containing 133-1034 bases of the rat OTR cDNA (NCBI Reference Sequence: NM_012871.3) subcloned into pSP73 Vector (Promega). The digoxigenin (DIG)-labeled antisense and sense RNA probe from the linearized *oxtr* cDNA template was synthesized using DIG RNA Labeling Kit (SP6/T7) (Roche Diagnostics). Sections containing 2 consecutive sections of the CeL (corresponding to Bregma: 2.5) were processed for fluorescent *in situ* hybridization (FISH). Rats were transcardially perfused with PBS followed by 4% PFA. Brains were dissected out and post fixed overnight in 4% PFA at 4 °C with gentle agitation. 50 µm vibratome sections were cut, collected and fixed in 4% PFA at 4°C overnight. The free-floating sections were washed in RNase-free PBS, immersed in 0.75% glycine in PBS, treated with 0.5 μg/ml proteinase K for 30 min at 37 °C, acetylated with 0.25% acetic anhydride in 0.1 M triethanolamine, and then hybridized with DIG-labeled RNA probe overnight at 65 °C. After RNase treatment and following intensive wash, the hybridized DIG-labeled probe was detected by incubation with Anti-Digoxigenin-POD (1:200; 11207733910; Roche Diagnostics) for 3 days at 4 °C. Signals were developed with tyramid signal amplification method. Rhodamine-conjugated tyramide was synthesized by coupling NHS-Rhodamine (Pierce Biotechnology, Thermo Fisher Scientific) to Tyramine-HCl (Sigma-Aldrich) in dimethylformamide with triethylamine. For the quantification of OTR mRNA-positive astrocytes, all confocal images were obtained using the same laser intensities and processed with the same brightness / contrast settings in Adobe Photoshop. Since the *in situ* signal for the OTR mRNA in astrocytes was weak, we first calculated the average intensity (signal intensity of all pixels divided by the total number of pixels) of the rhodamine-stained OTR mRNA signal for each individual section containing the CeL. Next, we calculated the standard deviation for each individual confocal image based on the intensity of all pixels comprising the image. We defined the threshold for OTR mRNA-positive astrocytes: If more than 1/4 of all pixels comprising an astrocyte soma displayed a signal intensity exceeding the average background intensity by more than 4-times the standard deviation, the astrocytes were considered as OTR mRNA-positive.

#### Astrocytes Markers

The aldehyde dehydrogenase 1 antibody is a commonly used marker for glial cells, including astrocytes. Therefore, we used the ALDH1L1 for immunohistochemistry in our initial experimental studies (Figure S1, S3). However, due to inconsistencies in staining quality as a result of batch-dependent antibody properties, especially in combination with the OTR mRNA FISH, we decided to employ Glutamine Synthetase (GluSyn, Figure 1a). GluSyn is a commonly used glial marker^71^, which stains astrocyte cell bodies, faint processes and even astrocytes not expressing GFAP. Using GluSyn, we achieved consistent results in combination with our OTR mRNA FISH.

#### Glutamine Synthase, ALDH1L1 Colocalization with OTR mRNA in Rat CeL

After development and washing steps the sections were stained with antibodies against glutamine synthase (mouse monoclonal, 1:500, ref: MAB302, MerckMilipore), ALDH1L1 (rabbit polyclonal, 1:500, ref: ab87117, abcam), in PBS and kept at 4°C on a shaker in a dark room overnight. After intensive washing with PBS, sections were stained with the respective secondary antibodies AlexaFluor488 (goat anti-mouse, 1:1000, ref: A11001, life technologies) and AlexaFluor680 (goat anti-mouse, 1:1000, ref: A27042, ThermoFischer Scientific) for 2 hours at RT. Following intensive washing with PBS, sections were mounted using Mowiol.

#### Double in situ Hybridizations for OTR mRNA and GFAP mRNA in Mice CeL

Fluorescent *in situ* hybridization (FISH) in Figure S1 was performed on 25-μm cryostat-cut coronal sections prepared from fresh-frozen mouse brain (male C57BL/6J, P22). After extraction, brains were immediately frozen in Tissue-Tek O.C.T. compound and stored at -80 degrees Celsius. ISH was performed according to the manufacturer’s instructions (Advanced Cell Diagnostics) for Fresh Frozen RNAscope Multiplex Fluorescent Assay. Treatment of amygdala containing sections were adjusted with the 3-plex negative control and then coexpression of OTR and GFAP examined using ACD designed target probes as well as the nuclear stain DAPI. Single plan images were collected with an upright laser scanning microscope (LSM-710, Carl Zeiss) using a 40x-objective with keeping acquisition parameters constant between control and probe treated sections.

#### Specific deletion of OTRs in CeL astrocytes

To specifically ablate OTRs in CeA astrocytes, transgenic cKO mice, in which *loxP* sites flank the OTR coding sequence (Lee et al., 2008), received bilateral injections (280 nl) of rAAV-GFAP-GFP-IRES-Cre. Following four weeks of expression of the viral proteins, mice were transcardially perfused with 1x PBS and 4% PFA. Brain sections were used for FISH (OTR mRNA) and IHC (GluSyn) to verify the validity of the approach. Representative images and quantifications are provided in Figure 2c.

#### AAV-Gfa-C1V1TSmCherry Specificity

After 3 weeks of vector expression in the brain, rats were transcardially perfused with 4% paraformaldehyde solution. Tissue blocks, containing CeA were dissected from the fixed brain and Vibratome-cut into 50 µm thick free-floating sections. After several rinse steps sampled sections were blocked with 5% NGS in PBS and incubated for 48 h at 4°C with polyclonal rabbit anti-ALDH1L1 antibody (1:500, Abcam) in 1% Triton-PBS buffer, containing 0,1 % NGS. Appropriate secondary antibody (AlexaFluor488 conjugated goat anti-rabbit (1:1000, LifeTechnologies) was used for further antigene detection. Intrinsic mCherry fluorescence of vector-expressing cells was strong enough to detect them in the tissue without any additional antibody enhancement. The immunolabeled sections were mounted onto Superfrost slides, cover-slipped with Moviol, analyzed and documented using LEICA SP5 confocal microscope.

#### Biocytin Filling of Astrocytes

In the lateral part of the central amygdala slices visualized with infrared-differential contrast optics, astrocytes were identified by their morphological appearance and the absence of action potentials in response to depolarizing current injections. Cells were patched with pipettes filled with (in mM) 110 K-gluconate, 30 KCl, 4 ATP-Mg, 10 phosphocreatine, 0.3 GTP, and 10 HEPES (pH: 7.3; 310 mOsm). Concentration of biocytin was 2 mg/ml. After obtaining whole-cell configuration astrocytes were hold at -80 mV and typical filling time was 30 minutes. Then the pipettes were carefully retracted and slices were incubated for additional 20 minutes in the oxygenated ACSF before fixation. Only one cell was filled per slice. Slices with filled cells were immersion-fixed at 4°C for 5 days in 4% PFA-PBS solution. Next, the slices were flat-embedded in 6% Agar-PBS, areas of interest were cut out of, re-embedded onto the Agar block and Vibratome-cut into 80 µm thick free-floating sections. The sections then were incubated with Avidin conjugated to Alexa Fluor488 (1:1000) (Thermo Fisher) in 1% Triton-PBS at 4°C, washed in PBS, mounted and cover-slipped. The tissue was analyzed and images taken at Leica TCS SP5 Confocal Microscope.

### Optogenetics

#### Ex vivo

To elicit evoked activation of OT axons in combination with CeL or CeM neurons patch-clamp, we used ChR2 expression in OT cells, as in^5^. Optical blue-light illumination of the CeL oxytocinergic axons expressing ChR2 was performed using light source XCite® 110LED from Excelitas Technologies through a GFP filter, controlled with Clampex driven TTL pulses for 20s at 30Hz with 10ms long pulses. To elicit light-evoked activation of OT axons in combination with astrocyte calcium imaging, or to elicit intracellular calcium increase in astrocytes, we used C1V1(t/t), a ChR1/VChR1 chimera with the combined mutations E122T/E162T (for more details, see^72^. Optogenetic green light stimulation of C1V1(t/t) in *ex vivo* experiments was performed using either the Spectra 7 LUMENCOR (λ542 nm) or light source X-Cite® 110LED from Excelitas Technologies through a Cy3 filter, controlled via MetaFluor or Clampex driven TTL pulses, respectively.

#### In vivo

Animals were habituated to the fixation of an optical fiber on the ferrule without light stimulation for one week before the experiment. In all cases, optical fibers were attached to the ferrules using an adapter (ADAF2, Thorlabs, NJ, USA) and animals let free to move in a typical home cage for the duration of the stimulation. Implanted optical fibers were connected to two lasers (LRS-0532-GFM-00100-03 LaserGlow 532nm DPSS Laser System) and the output power adjusted to correspond to 20 to 30 mW measured at the tip of 200 µm diameter fibers similar to the one implanted. Stimulation of 500 ms duration at a frequency of 0.5Hz were given for 3 min.

### Behavior

#### Mechanical Sensitivity Assessment

In experiments with rats, we used a calibrated forceps (Bioseb, Chaville, France) previously developed in our laboratory to test the animal mechanical sensitivity^73^. Briefly, the habituated rat was loosely restrained with a towel masking the eyes in order to limit stress by environmental stimulations. The tips of the forceps were placed at each side of the paw and a graduate force applied. The pressure producing a withdrawal of the paw, or in some rare cases vocalization, was considered as the nociceptive threshold value. This manipulation was performed three times for each hind paw and the values were averaged as being the final nociceptive threshold value. In experiments with mice, we used von Frey filaments tests. Mechanical allodynia (a symptom of neuropathic pain) was tested using von Frey hairs and results were expressed in grams. Tests were performed during the morning starting at least 2 h after lights on. Mice were placed in clear Plexiglas boxes (7 cm x 9 cm x 7 cm) on an elevated mesh floor. Calibrated von Frey filaments (Bioseb) were applied to the plantar surface of each hindpaw until they just bent in a series of ascending forces up to the mechanical threshold. Filaments were tested five times per paw and the paw withdrawal threshold (PWT) was defined as the lower of two consecutive filaments for which three or more withdrawals out of the five trials were observed.

#### Elevated Plus Maze

Following protocol from^74^, the arena is composed of four arms, two open (without walls) and two closed (with walls; rats 30 cm high; mice 15 cm high). Arms are 10 cm wide, 50 cm long and elevated 50 cm off the ground for rats and 5 cm wide, 30 cm long and elevetad 40 cm of the ground for mice. Two lamps with intensity adjustable up to 50 watts were positioned on the top of the maze, uniformly illuminating it. Animals were video tracked using a video-tracking systems (Ethovision Pro 3.16 Noldus, Wageningen, Netherlands and Anymaze, Stoelting Europe, Ireland). After each trial, the maze was cleaned with 70% ethanol and dry with paper towel. Twenty minutes after intracerebral injections or directly after optical stimulation, the animal was let free at the center of the plus maze, facing the open arm opposite to where the experimenter is, and was able to freely explore the entire apparatus for six minutes. Total time and time spend in closed and open arms were recorded in seconds and the percentage of time spent in closed arms was calculated as a measure of anxiety. As internal control, the total distance traveled during the test period was quantified and compared between all different groups (Figure S6a2-b2). Animals falling from the apparatus during the test, freezing more than 50% of the total time, or with cannulae/optic fiber issues, were removed from the analysis.

#### Conditioned Place Preference

The device is composed of two opaque conditioning boxes (rats: 30x32 cm; mice: 22x22 cm) and one clear neutral box (30x20 cm) Animals were video tracked using a video-tracking system (Anymaze, Stoelting Europe, Irland). After each trial, the device was cleaned with a disinfectant (Surfa’Safe, Anios laboratory). Based on^75^, all rats underwent a 3 days habituation period during which they were able to freely explore the entire apparatus for 30 min. On the day 3, behavior was record for 15min to verify the absence of pre-conditioning chamber preference. The time spend in the different compartment were measured and paired compartment was chosen as the compartment in which rat spent the less time during the 3rd day of habituation. On day 4, animals were placed the morning in one compartment for 15 min with no stimulation (unpaired box). Four hours after, the animal were placed 15min in the opposite box (paired box) and CeL astrocyte expressing C1V1 vector were optogenetically stimulated (3 min -500ms light pulse at 0.5 Hz - λ542nm) or TGOT micro-infused through intracerebral cannulae. On day 5, the animals were place in the CPP box and allowed to freely explore the entire apparatus during 15min. As internal control, the total distance traveled during the test period was quantified and compared between all different groups (Figure S6). Rats falling spending more than 80% of the total time in a single chamber before the conditioning, or with cannulae/optic fiber issues, were removed from the analysis.

## QUANTIFICATION AND STATISTICAL ANALYSIS

### Statistical Analysis

All parametrical statistical tests presented in figure captions were performed following correct verification of the assumptions on the distribution of data, and if not non-parametric tests were used. Tests were performed using either SPSS 23 (IBM) or statview 5 (SAS institute Inc.). All values, group compositions and statistical tests for each experiment and figure panel are detailed in Supplementary Tables 1-6.

### Spatial Distribution Analysis of Astrocytes within the CeL

For the analysis of spatial distribution of astrocytes within the CeL, confocal images were analyzed and all OTR+ astrocytes were manually marked. A separate image with the location of identified astrocytes in the CeL was exported for spatial distribution analysis. The coordinate’s points of OTR+ astrocytes (events) were extracted with custom-written MATLAB (MathWorks) script and the centroid of each event was calculated with the function *regionprops*. The univariate first-order nearest neighbor (NN) test (Clark-Evans statistic) was derived from the distance to the closest event for each data point in the image. The test estimates the departure of the observed NN distances from those expected under a null model of complete spatial randomness (CSR; a homogeneous Poisson process). We calculated the mean NN distance for all events in each image as:

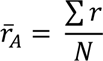

where *r* is the distance from a given event to its nearest neighbor, and *N* is the number of events in a given population. The ratio 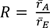 is the observed mean NN distance by mean distance which would be expected if this population was distributed randomly. In a random distribution *R* = 1. The values of *R* can vary between 0 (maximum aggregation or *clustering*) and 2.1491 (maximum spacing or *uniformity*). For details see^76, 77^. Thus, *R* values are indicative of uniform, random, and clustered patterns of point distribution. To evaluate the significance of *R*, it is possible to calculate the standard variate of the normal curve, namely the c-score:

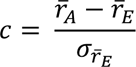

where 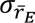 is the standard error of the mean distance to NN in a randomly distributed population with the same number of events as the observed population. The c value 1.96 represents the 5% level of significance for a two-tailed test. We calculated the mean NN distance for 12 randomly selected images, *R*, and the c-score for image of the CeL (Figure S1b): Only in two images the c-score was equal to or greater than 1.96, therefore only in two of the images it was possible to reject the null hypothesis of randomly distributed population (Statistics in Table S1).

## SUPPLEMENTAL INFORMATION

**Figure S1.**
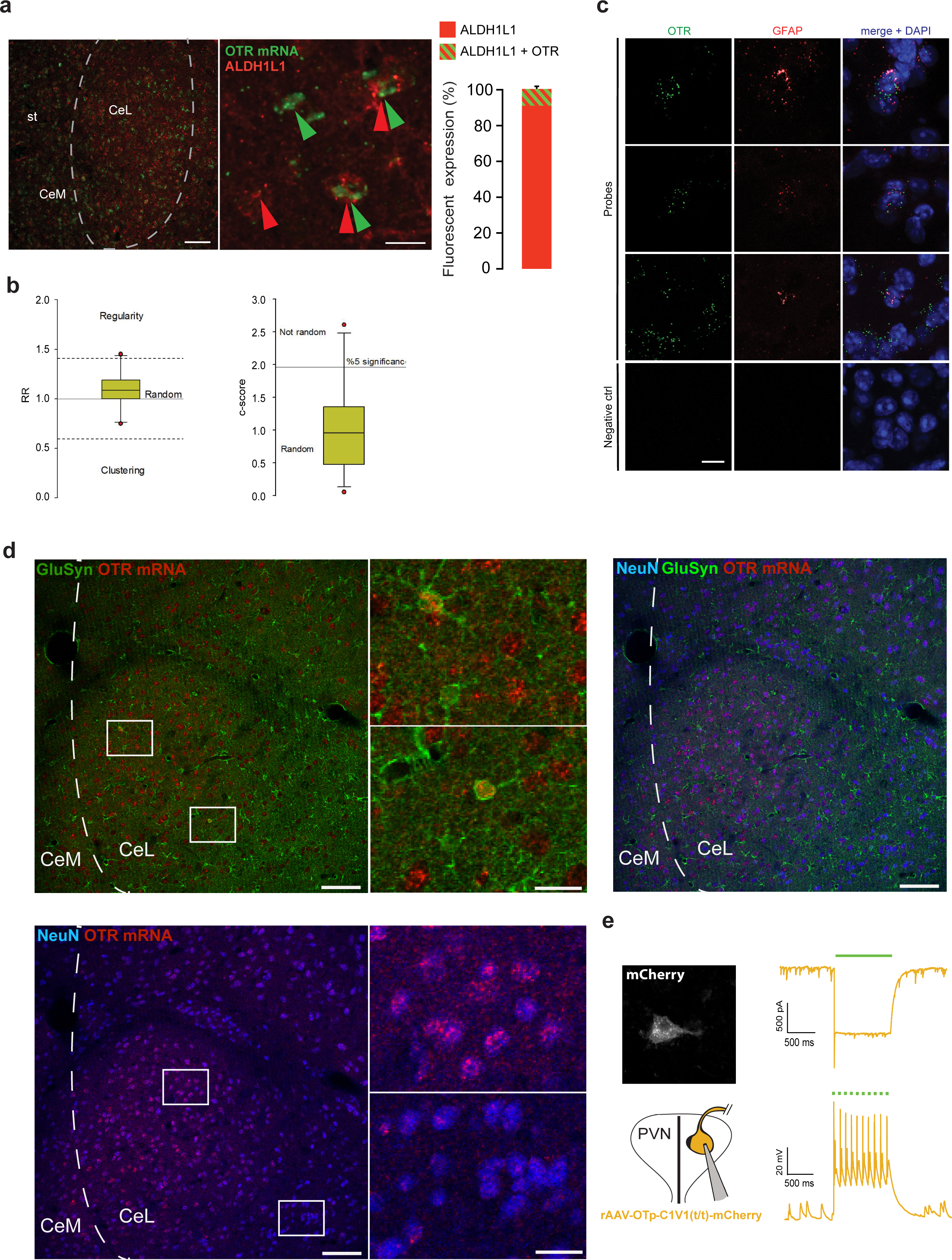
(**a**) (Left) FISH overview of CeA hybridized with OTR mRNA (green), counterstained with polyclonal anti-ALDH1L1 antibody (red). (Middle) High magnification image of cells positive for both OTR mRNA and ALDH1L1 (double arrows); green arrows point OTR mRNA-positive cells; red arrows point ALDH1L1-positive cells. Scale bar 400 (Left) and 50 µm (Middle). (Right) Quantification of ALDH1L1-positive cells containing OTR mRNA signaling. n astrocytes=450; n rats=3 (**b**) Spatial distribution analysis of astrocytes within the CeL (R and c-score) demonstrating the random distribution of OTR positive astrocytes. (**c**) RNAscope *in situ* hybridization showing GFAP (red) and OTR (green) expressing cells in mice CeA. Merged images include DAPI stain (blue). (Bottom) Negative control probe targeting the bacterial gene DapB. Scale bar 10μm. (**d**) Combination of FISH and IHC showing co-localization of GluSyn (top left), NeuN (bottom) and both markers (top right) with OTR mRNA in mice. 1254 NeuN-positive neurons and 1185 GluSyn-positive astrocytes. We detected 67.8 % OTR-positive neurons, mostly in the dorsal part the the CeL, and 12.7 % OTR-positive astrocytes. Scale bars 100µm, 10µm. (**e**) (Left) C1V1(t/t)-mCherry expressing oxytocin neurons of the PVN in which (Right) λ542nm light exposure induced depolarizing currents, enabling precise spiking control as measured in whole cell patch-clamp. st: stria terminalis. (Statistics and numbers in Table S1).

**Figure S2.**
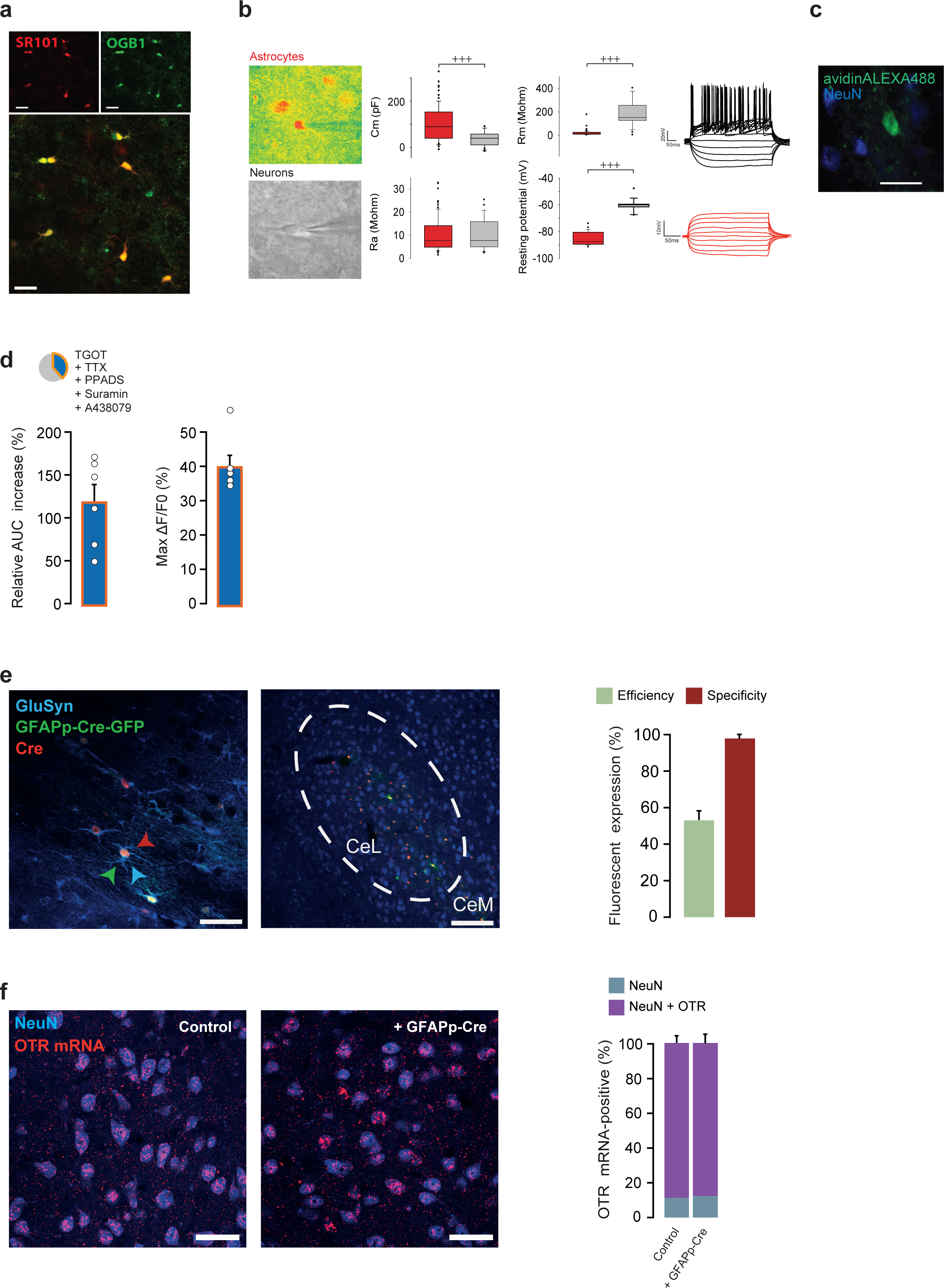
(**a**) Typical confocal image of CeL astrocytes co-labeled with SR101 and OGB1. Scale bar 20μm. (**b**) (Left) Pseudo-color pictures of an SR101 positive cell identified as an astrocyte compared to neurons identified under oblique infrared light, Scale bar 10μm. (Middle) Electrophysiological properties of patched SR101+ (red, n=84) and SR101-cells (grey, n=21). (Right) Typical responses to 20pA current steps of a SR101+ (red) and a SR101-cells (black) (**c**) CeL SR101 positive cell filled with biocytin through whole cell patch-clamp (green) lacks NeuN signal (Blue). Scale bar 50 μm. (**d)** (Up) Pie charts of the proportion of TGOT+TTX responsive astrocytes and their (bottom, histograms) relative increase of ΔF/F AUC after drugs application and maximal peak values upon exposition to PPADS (50 µM) + Suramin (75 µM) + A438079 (1 µM) (ns = 6, na = 289). Data are expressed as mean across slices plus SEM. White circles indicate average across responding astrocytes per slice. \**P*<0.05, independent samples Students’s t-test, ^+^*P*<0.05, ^++^*P*<0.01, ^+++^*P*<0.001 independent samples Mann-Whitney U test, ^#^*P*<0.05 Wilcoxon signed rank test (Statistics in Table S2). White circles represent the average across responding astrocytes per slice. (**e**) Immunohistochemical staining for glutamine synthetase (GluSyn; *i.e*. astrocytes; blue), GFP (green) and Cre recombinase (red). (Right picture) Overview of the CeA, displaying correct viral targeting of the CeL subdivision. Efficiency: Over 1001 GluSyn positive cells, 561 were also GFP positive, indicating an efficiency of 56±4.9%). Specificity: we counted a total of n=977 GFP-positive cells, 940 of which were positive for GluSyn (96.2±2.1%). None of the GFP or Cre signals were detected in NeuN positive cells (0 out of n = 850, 3 mice, not shown). Finally, 99.82 ± 0.2% of GluSyn-positive astrocytes contained GFP signal were Cre-positive (n = 1001). (**f**) Immunohistochemical analysis of OTR expression in NeuN-positive neurons of the CeL revealed no difference in OTR levels between control and GFAP-Cre injected animals; 89.3 ± 4.8% (n = 989, n = 4 mice) and 86.0 ± 3.7% (n = 1204, n = 4 mice) respectively. (Statistics and numbers in Table S2).

**Figure S3.**
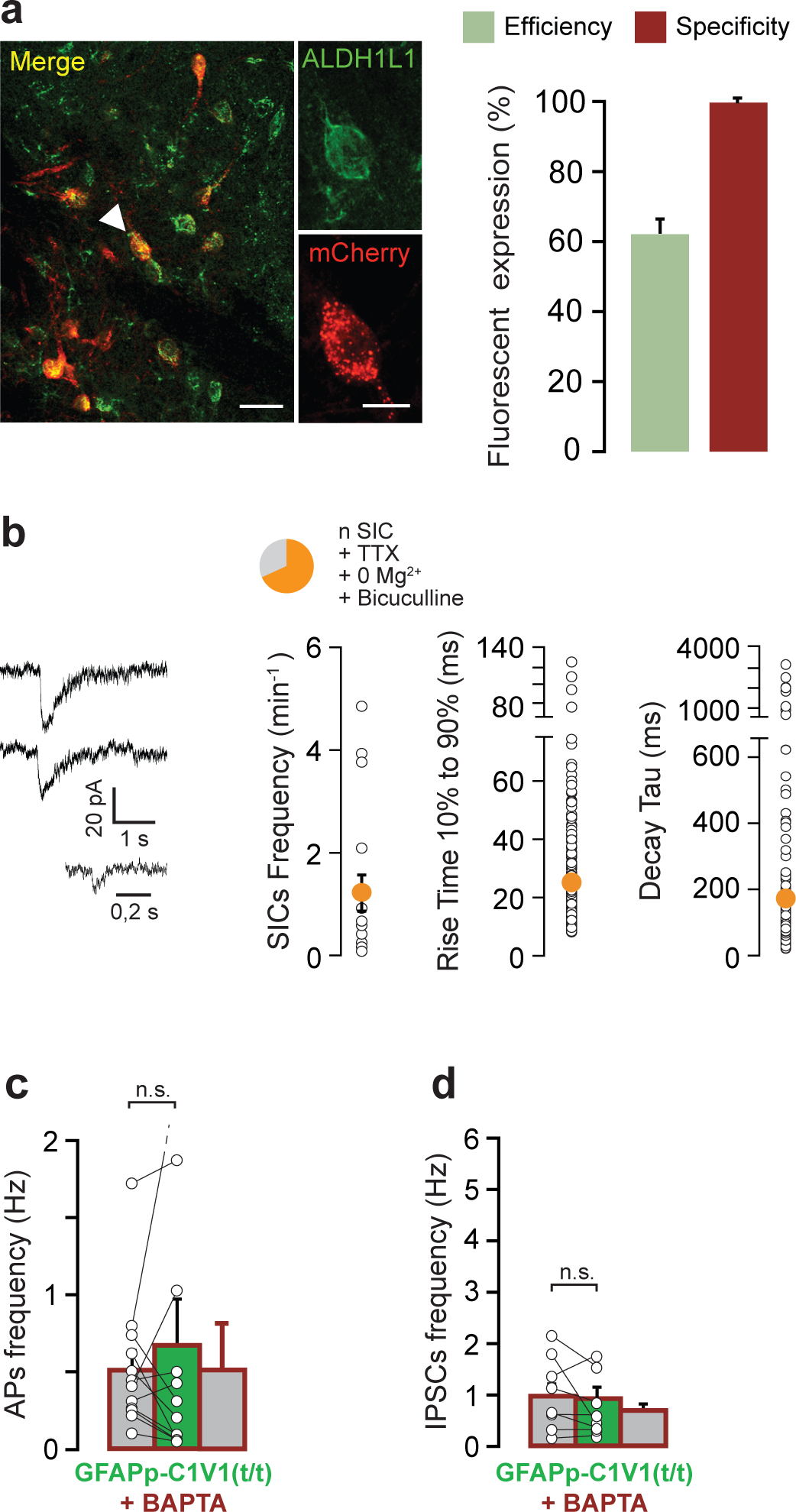
(**a**) (Left) Immunohistochemistry image shows CeL cells transfected with rAAV-Gfap-C1V1TSmCherry with co-labeling for ALDH1L1. White arrow shows one cell expanded in insets. Scale bars 25 and 10 µm (insets). (Right) Quantification of the efficiency and specificity of transduction of C1V1 in CeL astrocytes. rAAV GFAP-C1V1-mCherry vector was injected into rat CeA (bilaterally, 300nl). Specificity: we counted a total of n=323 mCherry-positive cells, 318 of which were positive for ALDH1L1 (98.5±0.8%). None of the analyzed cells were positive for NeuN. Efficiency: Over 517 ALDH1L1 positive cells, 318 were also mCherry positive, indicating an efficiency of 62.3±3.5%). (**b**) (Top) Pie chart showing the proportion CeL neurons displaying SICs and (bottom) their electrophysiological properties, measured in 0 Mg^2+^ ACSF with TTX (1 µM) and Bicuculline (10 µM) (n = 15). (**c**, **d**) Effect of CeL astrocytes BAPTA loading on 3 min long light-evoked activation of C1V1 expressing CeL astrocytes (1 s width, 0.5 Hz) on APs (**c**) and IPSCs (**d**) frequencies recorded in CeL and CeM neurons (n = 12 and 9, respectively). In (b, c, d) data are expressed as averages plus SEM and white circles represents SICs electrophysiological properties averaged per cell. (Statistics and numbers in Table S3).

**Figure S4.**
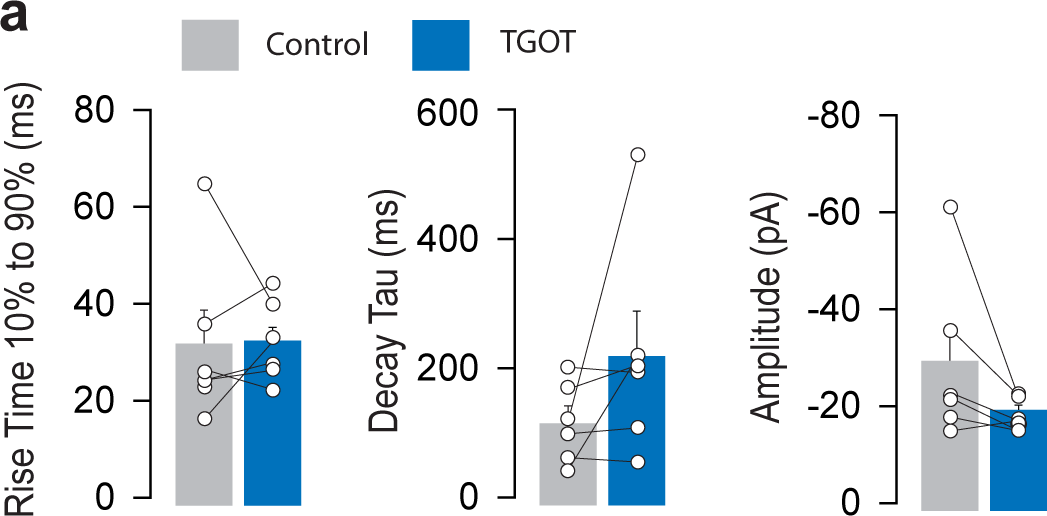
(**a**) Effect of TGOT on SICs properties in presence of 0 Mg^2+^, TTX (1µM) and Bicuculline (10 µM) (n = 6). Data are expressed as averages plus SEM and white circles represents SICs electrophysiological properties averaged per cell before and after TGOT application. (Statistics and numbers in Table S4).

**Figure S5.**
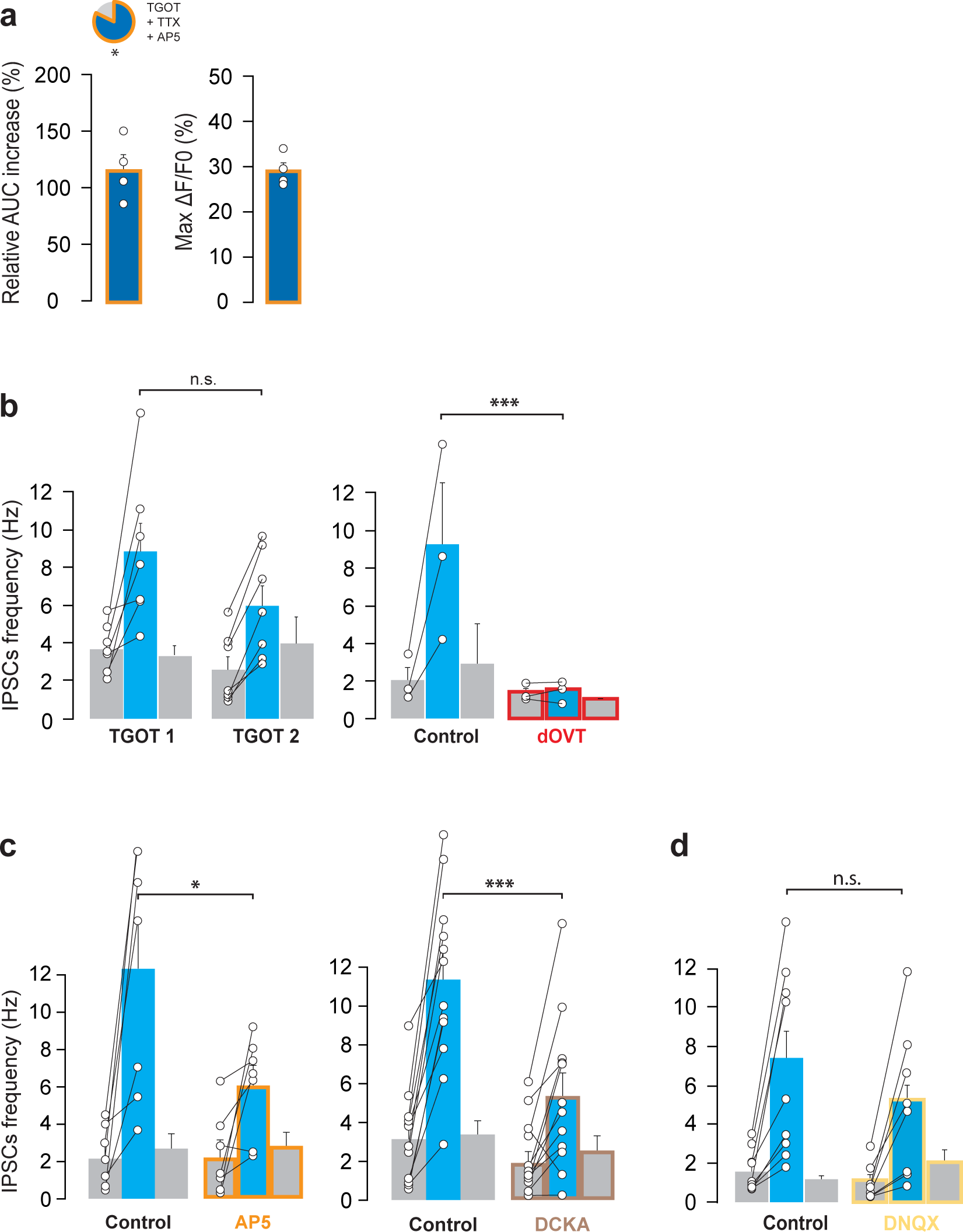
(**a**) (Top) pie chart of the proportion of TGOT+TTX responsive astrocytes and (bottom) relative increase of ΔF/F AUC after TGOT application and maximal peak value following exposition to AP5 (50 µM; ns = 4, na = 35). (**b**) (Left) Effect of double (20 min apart) application of TGOT on IPSCs frequencies in CeM (0.4 µM n = 7). (Right) dOVT (1 µM; n = 3) prevents the effect of TGOT on IPSCs frequencies in CeM. (**c**) Effect of AP5 (50 µM, n = 7) or DCKA (10 µM, n = 12) on TGOT-induced increase in IPSCs frequencies in CeM neurons. (**d**) DNQX (25 µM; n = 8) effect on TGOT induced increase in IPSCs frequencies in CeM neurons. In (b,c,d) Data are expressed as averages plus SEM and white circles represent individual cell data. * *P*<0.05, ** *P*<0.01, independent samples (calcium-imaging) or paired (patch-clamp) Students’s t-test (Statistics and numbers in Table S5).

**Figure S6.**
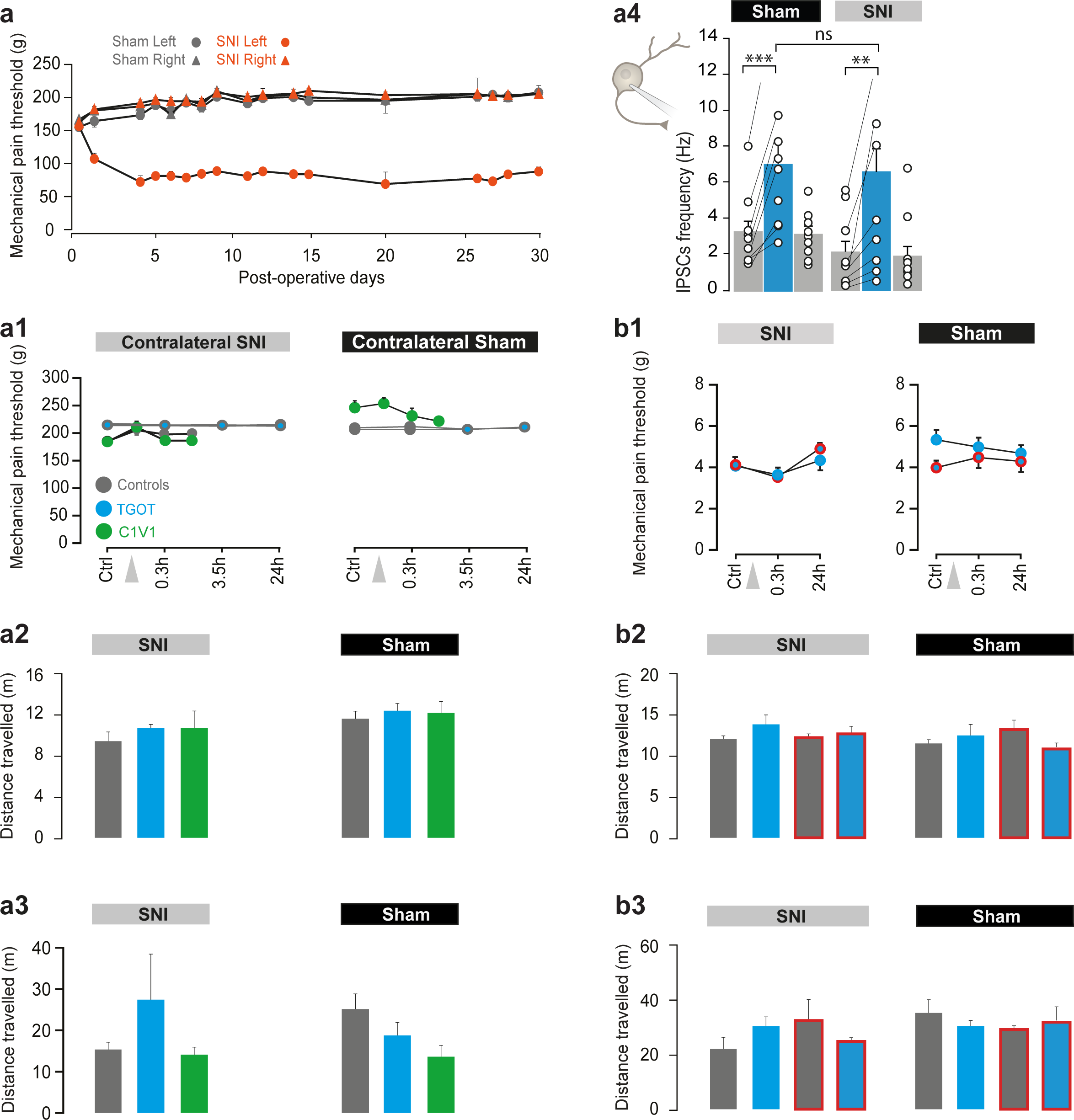
(**a**) Experiments in rat model. 30 days post surgeries time course of mechanical pain threshold evolution across sham and SNI groups. (**b**) Experiments in mice model. (**a1, b1**) Mechanical pain threshold was also assessed on the non-injured paw of SNI (left) and its equivalent in Sham (right) groups. TGOT or its vehicle, or astrocytes light-evoked activation of C1V1, were administered in the CeL and mechanical pain threshold assessed again at different time points. (**a2, b2**) Locomotion was assessed through measurement of the distance travelled during the length of the elevated plus maze experiment, after administration of the different treatments. (**a3, b3**) Locomotion was assessed through measurement of the distance travelled during the time of the conditioned place preference experiment, after administration of the different treatments. (**a4**) TGOT effect on CeM neurons IPSCs frequencies is unchanged between Sham (n = 16) or SNI (n = 16) rats. Data are expressed as average plus SEM and white circles represent individual cell data. Mixed design ANOVA followed by posthoc Bonferroni tests (Statistics in Table S6).

**Table S1.** Numerical Values and Statistical Analysis of Data presented in **Fig. 1**, and **S1**

| Figure | Statistical unit | Groups tested | Paired or unpaired | N | Unit | Mean | SEM |
| --- | --- | --- | --- | --- | --- | --- | --- |
| 1c | Proportion of rat OTR positive astrocytes on 16 sections |  |  | 1185 | % | 12,70 | 3,43587083 |
|  | Proportion of rat OTR positive neurons on 16 sections |  |  | 1254 | % | 67,80 | 3,17608249 |
| 1g | onset time of responses | CeL astrocytes |  | 21 | s | 42,500 | 15,853 |
|  | averages of offset time of responses | CeL astrocytes |  | 21 | s | 435,762 | 67,643 |
| S1a | Proportion of OTR positive |  |  | 450 | % | 8,67 | 1,07 |

| Figure | Statistical unit | Image | N | $r_A^-$ [μm] | R | c-score |
| --- | --- | --- | --- | --- | --- | --- |
| S1b | Spatial distribution analysis of astrocytes within the CeL | Image 1 | 6 | 92 ± 22 | 0,75 | 1,16 |
| | | Image 2 | 8 | 116 ± 15 | 1,1 | 0,52 |
| | | Image 3 | 10 | 103 ± 9 | 1,03 | 0,49 |
| | | Image 4 | 12 | 100 ± 21 | 1,16 | 1,04 |
| | | Image 5 | 10 | 100 ± 23 | 1,05 | 0,33 |
| | | Image 6 | 12 | 104 ± 13 | 1,2 | 1,3 |
| | | Image 7 | 12 | 93 ± 13 | 1,07 | 0,47 |
| | | Image 8 | 12 | 69 ± 20 | 0,79 | 1,37 |
| | | Image 9 | 12 | 99 ± 26 | 0,99 | 0,05 |
| | | Image 10 | 13 | 94 ± 11 | 1,13 | 0,87 |
| | | Image 11 | 8 | 149 ± 43 | 1,4 | 2,18 |
| | | Image 12 | 9 | 145 ± 32 | 1,45 | 2,61 |

**Table S2.**
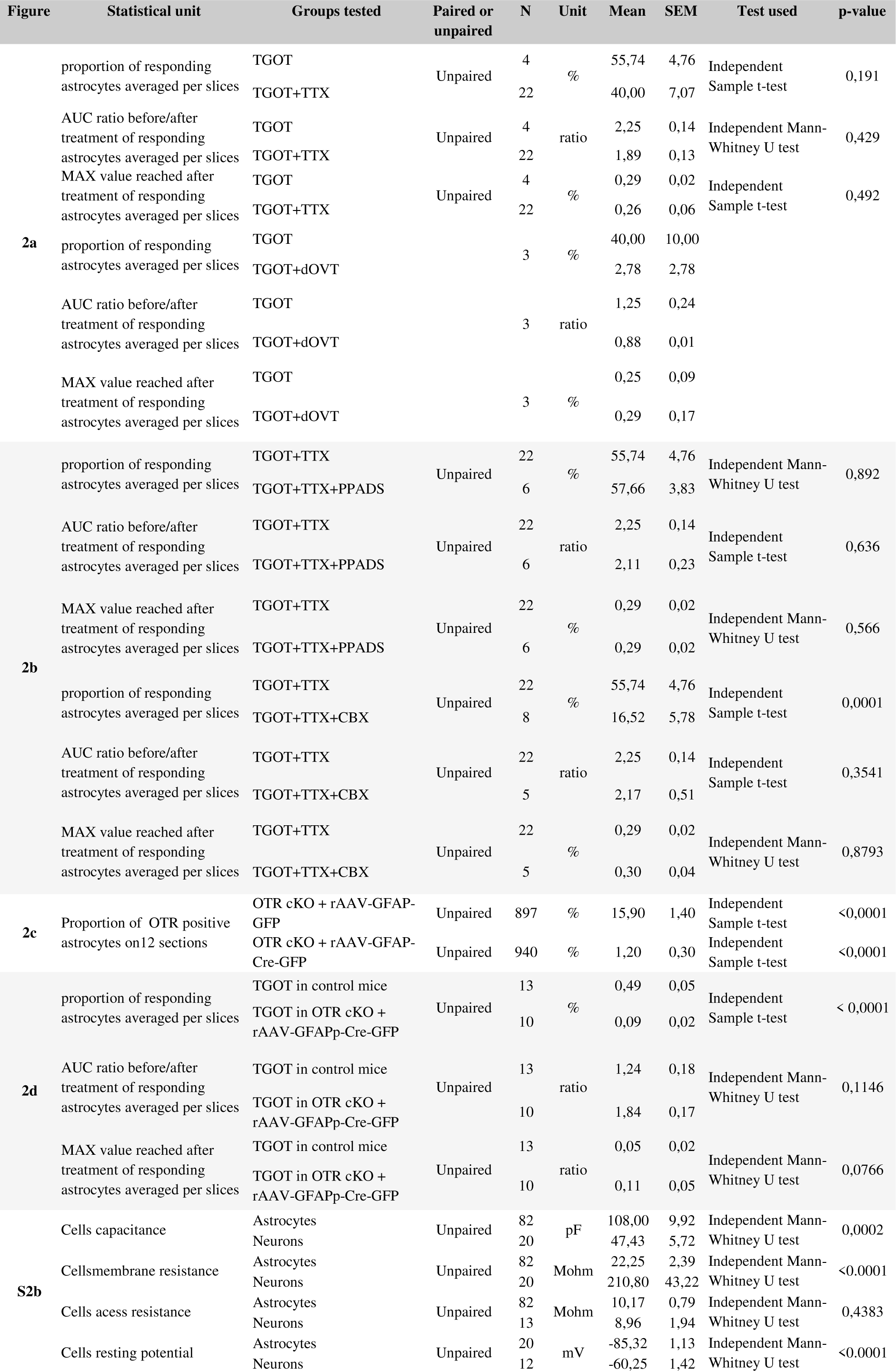
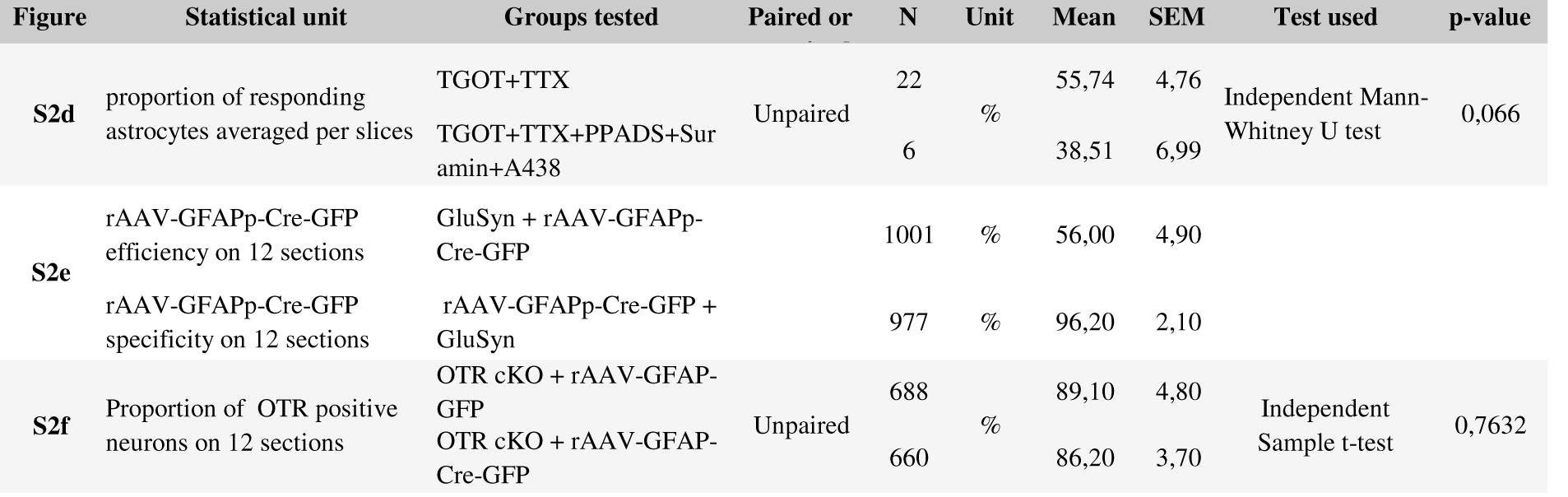
Numerical Values and Statistical Analysis of Data presented in **Fig. 2** and **S2**

**Table S3.** Numerical Values and Statistical Analysis of Data presented in **Fig. 3** and **S3**

| Figure | Statistical unit | Groups tested | Paired or unpaired | N | Unit | Mean | SEM | Test used | p-value |
| --- | --- | --- | --- | --- | --- | --- | --- | --- | --- |
| **3c** | SICs frequencies across neurons recorded before and after treatment | Baseline | Paired | 19 | Hz | 0,8457 | 0,2363 | Wilcoxon signed rank test | 0,0064 |
| | | 20s λ542nm light pulse | | | | 1,517 | 0,3781 | | |
| **3d** | APs frequencies across neurons recorded before and after treatment | Baseline | Paired | 10 | Hz | 0,25 | 0,1256 | Dunn's multiple comparisons | 0,0003 |
| | | 20s λ542nm light pulse | | | | 0,745 | 0,3044 | | |
|  |  | Wash |  |  |  | 0,3 | 0,1287 |  |  |
| **3e** | IPSCs frequencies across neurons recorded before and after treatment | Baseline | Paired | 9 | Hz | 1,955 | 0,358 | Wilcoxon signed rank test | 0,0038 |
| | | 20s λ542nm light pulse | | | | 3,368 | 0,635 | | |
| **S3a** | rAAV-GFAPp-C1V1-mCherry specificity on 9 sections | rAAV-GFAPp-C1V1-mCherry + ALDH1L1 |  | 323 | % | 98,452 | 0,802 |  |  |
|  |  | rAAV-GFAPp-C1V1-mCherry + NeuN |  | 318 | % | 0,000 | 0,000 |  |  |
|  |  | ALDH1L1 + rAAV-GFAPp-C1V1-mCherry |  | 517 | % | 62,300 | 3,500 |  |  |
| **S3c** | APs frequencies across neurons recorded before and after treatment | Baseline+BAPTA | Paired | 12 | Hz | 0,5292 | 0,1274 | Dunn's multiple comparisons test | 0,0395 |
| | | 20s λ542nm light pulse+BAPTA | | 12 | | 0,6833 | 0,3054 | | |
| **S3d** | IPSCs frequencies across neurons recorded before and after treatment | Baseline+BAPTA | Paired | 9 | Hz | 0,994 | 0,222 | Holm-Sidak's multiple comparisons test | 0,072 |
| | | 20s λ542nm light pulse+BAPTA | | 9 | | 0,992 | 0,197 | | |

**Table S4.** Numerical Values and Statistical Analysis of Data presented in **Fig. 4** and **S4**

| Figure | Statistical unit | Groups tested | Paired or unpaired | N | Unit | Mean | SEM | Test used | p-value |
| --- | --- | --- | --- | --- | --- | --- | --- | --- | --- |
| **4a1** | SICs frequencies under TGOT across neurons recorded before and after treatment | Baseline | Paired | 7 | Hz | 1,10 | 0,5088 | Wilcoxon signed rank test | 0,0156 |
|  |  | TGOT |  | 7 |  | 2,50 | 0,6437 |  |  |
|  |  | Baseline+Bapta | Paired | 8 | Hz | 0,90 | 0,2478 | Wilcoxon signed rank test | > 0,9999 |
|  |  | Bapta +TGOT |  | 8 |  | 0,88 | 0,2698 |  |  |
|  |  | TGOT | Unpaired | 7 | Hz | 2,50 | 0,6437 | Independent Mann-Whitney U test | 0,0197 |
|  |  | Bapta +TGOT |  | 8 |  | 0,88 | 0,2698 |  |  |
| **4a2** | APs frequencies under TGOT across neurons recorded before and after treatment | Baseline | Paired | 9 | Hz | 0,43 | 0,1277 | Wilcoxon signed rank test | 0,0039 |
|  |  | TGOT |  | 9 |  | 0,92 | 0,1732 |  |  |
|  |  | Baseline+BAPTA | Paired | 9 | Hz | 0,20 | 0,0898 | Wilcoxon signed rank test | 0,0938 |
|  |  | Bapta +TGOT |  | 9 |  | 0,12 | 0,0595 |  |  |
|  |  | TGOT | Unpaired | 9 | Hz | 0,92 | 0,1732 | Independent Mann-Whitney U test | 0,0005 |
|  |  | Bapta +TGOT |  | 9 |  | 0,12 | 0,0595 |  |  |
| **4a3** | IPSCs frequencies under TGOT across neurons recorded before and after treatment | TGOT | Unpaired | 17 | Hz | 5,05 | 0,691 | Independent Mann-Whitney U test | p<0,001 |
|  |  | Bapta +TGOT |  | 17 |  | 1,31 | 0,351 |  |  |
| **4b1** | SICs frequencies under TGOT across neurons recorded before and after treatment | baseline in control mice | paired | 8 | Hz | 1,10 | 0,3397 | Wilcoxon signed rank test | 0,0391 |
|  |  | TGOT in control mice |  | 8 |  | 1,71 | 0,3669 |  |  |
|  |  | Baseline in OTR cKO mice + rAAV-Cre-GFP | Paired | 5 | Hz | 0,76 | 0,1939 | Wilcoxon signed rank test | > 0,9999 |
|  |  | TGOT in OTR cKO mice + rAAV-Cre-GFP |  | 5 |  | 0,75 | 0,1921 |  |  |
|  |  | TGOT in control mice | Unpaired | 8 | Hz | 1,71 | 0,3669 | Independent Mann-Whitney U test | 0,0862 |
|  |  | TGOT in OTR cKO mice + rAAV-Cre-GFP |  | 5 |  | 0,75 | 0,1921 |  |  |
| **4b2** | APs frequencies under TGOT across neurons recorded before and after treatment | baseline in control mice | Paired | 7 | Hz | 0,26 | 0,1071 | Wilcoxon signed rank test | 0,0156 |
|  |  | TGOT in control mice |  | 7 |  | 0,82 | 0,1495 |  |  |
|  |  | Baseline in OTR cKO mice + rAAV-Cre-GFP | Paired | 11 | Hz | 0,35 | 0,0691 | Wilcoxon signed rank test | 0,8203 |
|  |  | TGOT in OTR cKO mice + rAAV-Cre-GFP |  | 11 |  | 0,40 | 0,1263 |  |  |
|  |  | TGOT in control mice | Unpaired | 7 | Hz | 0,82 | 0,1495 | Independent Mann-Whitney U test | 0,032 |
|  |  | TGOT in OTR cKO mice + rAAV-Cre-GFP |  | 11 |  | 0,40 | 0,1263 |  |  |
| **4b3** | IPSCs frequencies under TGOT across neurons recorded before and after treatment | Baseline in control mice | Paired | 27 | Hz | 1,52 | 0,2621 | Wilcoxon signed rank test | < 0,0001 |
|  |  | TGOT in control mice |  | 27 |  | 2,80 | 0,3876 |  |  |
|  |  | Baseline in OTR cKO mice + rAAV-Cre-GFP | Paired | 16 | Hz | 1,72 | 0,3677 | Wilcoxon signed rank test | 0,9501 |
|  |  | TGOT in OTR cKO mice + rAAV-Cre-GFP |  | 16 |  | 1,93 | 0,4365 |  |  |
|  |  | TGOT in control mice | Unpaired | 27 | Hz | 1,52 | 0,2621 | Independent Mann-Whitney U test | 0,0353 |
|  |  | TGOT in OTR cKO mice + rAAV-Cre-GFP |  | 16 |  | 1,93 | 0,4365 |  |  |
| Figure | Statistical unit | Groups tested | Paired or unpaired | N | Unit | Mean | SEM | Test used | p-value |
| **S3c** | 10 to 90% rise time of SICs across cells before and after treatment | Baseline | Paired | 6 | ms | 31,68 | 7,158 | Paired t-test | 0,922 |
|  |  | TGOT |  |  |  | 32,28 | 3,482 |  |  |
|  | decay Tau of SICs across cells before and after treatment | Baseline | Paired | 6 | ms | 113,78 | 25,468 | Wilcoxon signed rank test | 0,116 |
|  |  | TGOT |  |  |  | 217,26 | 68,062 |  |  |
|  | amplitude of SICs across cells before and after treatment | Baseline | Paired | 6 | pA | -27,79 | 7,134 | Paired t-test | 0,141 |
|  |  | TGOT |  |  |  | -17,22 | 1,304 |  |  |

**Table S5.** Numerical Values and Statistical Analysis of Data presented in **Fig. 5** and **S5**

| Figure | Statistical unit | Groups tested | Paired or unpaired | N | Unit | Mean | SEM | Test used | p-value |
| --- | --- | --- | --- | --- | --- | --- | --- | --- | --- |
| 5c | SICs frequencies across CeL neuron before and after treatment | Baseline | Paired | 7 | n/min | 1,100 | 0,509 | Wilcoxon signed rank test | 0,0156 |
|  |  | TGOT |  | 7 |  | 2,500 | 0,644 |  |  |
|  |  | Ifenprodil | Paired | 10 |  | 0,425 | 0,084 | Wilcoxon signed rank test | 0,5703 |
|  |  | Ifenprodil + TGOT |  | 10 |  | 0,367 | 0,136 |  |  |
|  |  | Baseline | Unpaired | 7 |  | 1,100 | 0,509 | Independent Mann-Whitney U test | 0,0409 |
|  |  | AP5 |  | 10 |  | 0,105 | 0,027 |  |  |
| 5d | APs frequencies across CeL neuron before and after treatment | Baseline | Paired | 9 | Hz | 0,433 | 0,128 | Wilcoxon signed rank test | 0,0039 |
|  |  | TGOT |  | 9 |  | 0,917 | 0,173 |  |  |
|  |  | Baseline AP5 | Paired | 7 | Hz | 0,464 | 0,108 | Wilcoxon signed rank test | 0,5313 |
|  |  | TGOT + AP5 |  | 7 |  | 0,407 | 0,150 |  |  |
|  |  | TGOT | Unpaired | 9 | Hz | 0,917 | 0,173 | Mann Whitney test | 0,0122 |
|  |  | AP5 + TGOT |  | 7 |  | 0,407 | 0,150 |  |  |
| 5e | IPSC frequencies across neurons recorded before and after | TGOT+DAAO | Paired | 9 | Hz | 0,611 | 0,182 | Wilcoxon signed rank test | 0,0076 |
|  |  | TGOT+D-Serine |  |  |  | 2,711 | 0,634 |  |  |
| Figure | Statistical unit | Groups tested | Paired or unpaired | N | Unit | Mean | SEM | Test used | p-value |
| S5a | proportion of responding astrocytes averaged per slices | TGOT+TTX | Unpaired | 22 | % | 55,739 | 4,764 | Independent Sample t-test | 0,037 |
|  |  | TGOT+TTX+AP5 |  | 4 |  | 82,292 | 10,259 |  |  |
|  | AUC ratio before/after treatment of responding astrocytes averaged per slices | TGOT+TTX | Unpaired | 22 | ratio | 2,253 | 0,142 | Independent Sample t-test | 0,795 |
|  |  | TGOT+TTX+AP5 |  | 4 |  | 2,163 | 0,138 |  |  |
|  | MAX value reached after treatment of responding astrocytes averaged per slices | TGOT+TTX | Unpaired | 22 | % | 0,293 | 0,021 | Independent Sample t-test | 0,941 |
|  |  | TGOT+TTX+AP5 |  | 4 |  | 0,290 | 0,018 |  |  |
| S5b | IPSCs frequencies under TGOT across neurons recorded before and after treatment | TGOT1 | Paired | 7 | Hz | 8,900 | 1,500 | Paired t-test | 0,0529 |
|  |  | TGOT2 |  |  |  | 6,007 | 1,062 |  |  |
|  | IPSC frequencies under TGOT across neurons recorded before and after treatment | TGOT |  | 3 | Hz | 9,217 | 3,028 |  |  |
|  |  | TGOT+dOVT |  |  |  | 1,383 | 0,328 |  |  |
| S5c | IPSC frequencies under TGOT across neurons recorded before and after treatment | TGOT | Paired | 7 | Hz | 12,371 | 2,558 | Paired t-test | 0,0308 |
|  |  | TGOT+AP5 |  |  |  | 6,121 | 1,018 |  |  |
|  | IPSC frequencies under TGOT across neurons recorded before and after treatment | TGOT | Paired | 12 | Hz | 11,300 | 1,336 | Paired t-test | <0.001 |
|  |  | TGOT+DCKA |  |  |  | 5,338 | 1,133 |  |  |
| S5d | IPSC frequencies under TGOT across neurons recorded before and after treatment | TGOT | Paired | 8 | Hz | 7,316 | 1,811 | Dunn's multiple comparisons | >0.05 |
|  |  | TGOT+DNQX |  |  |  | 5,268 | 1,395 |  |  |

**Table S6.**
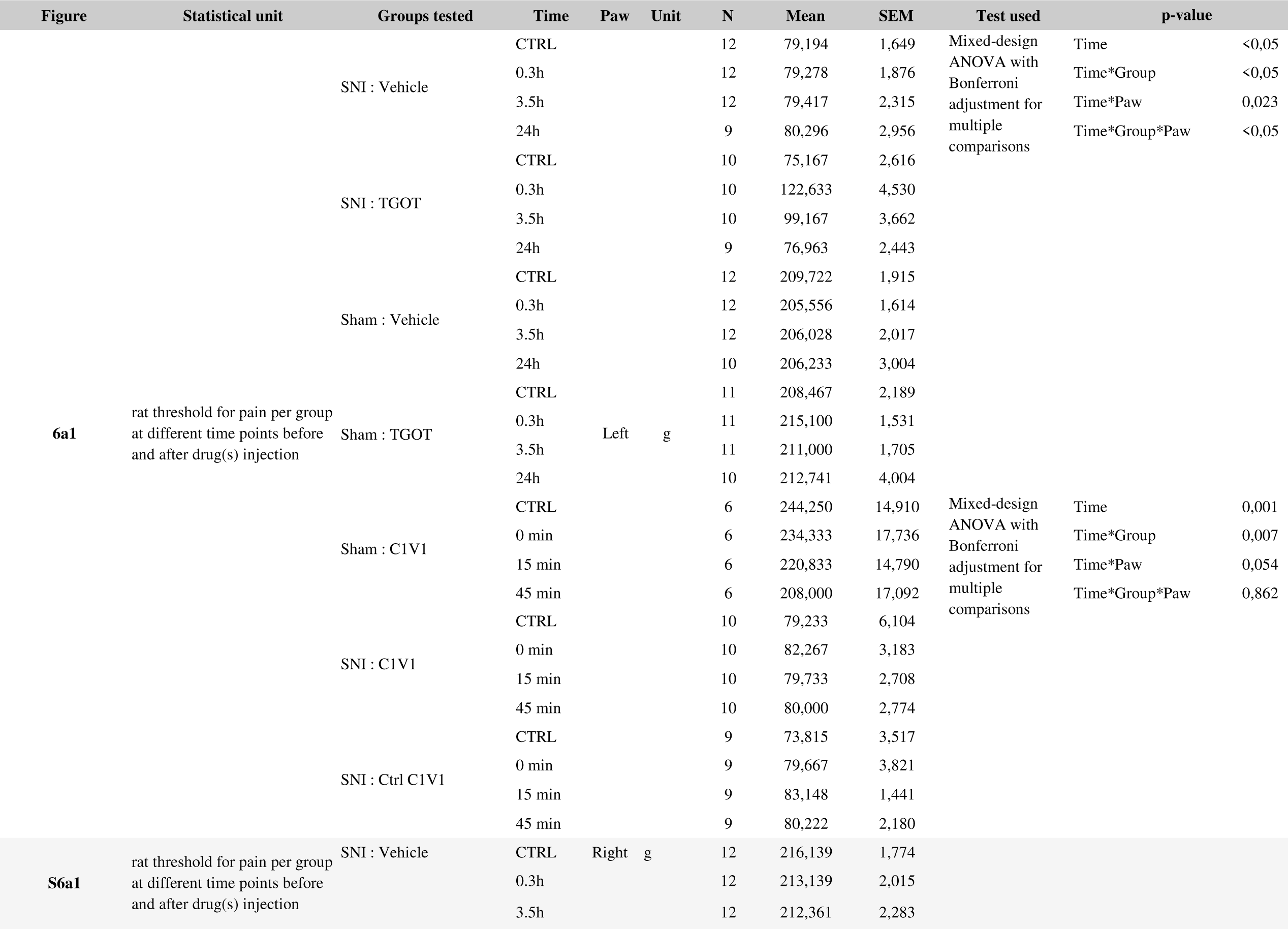
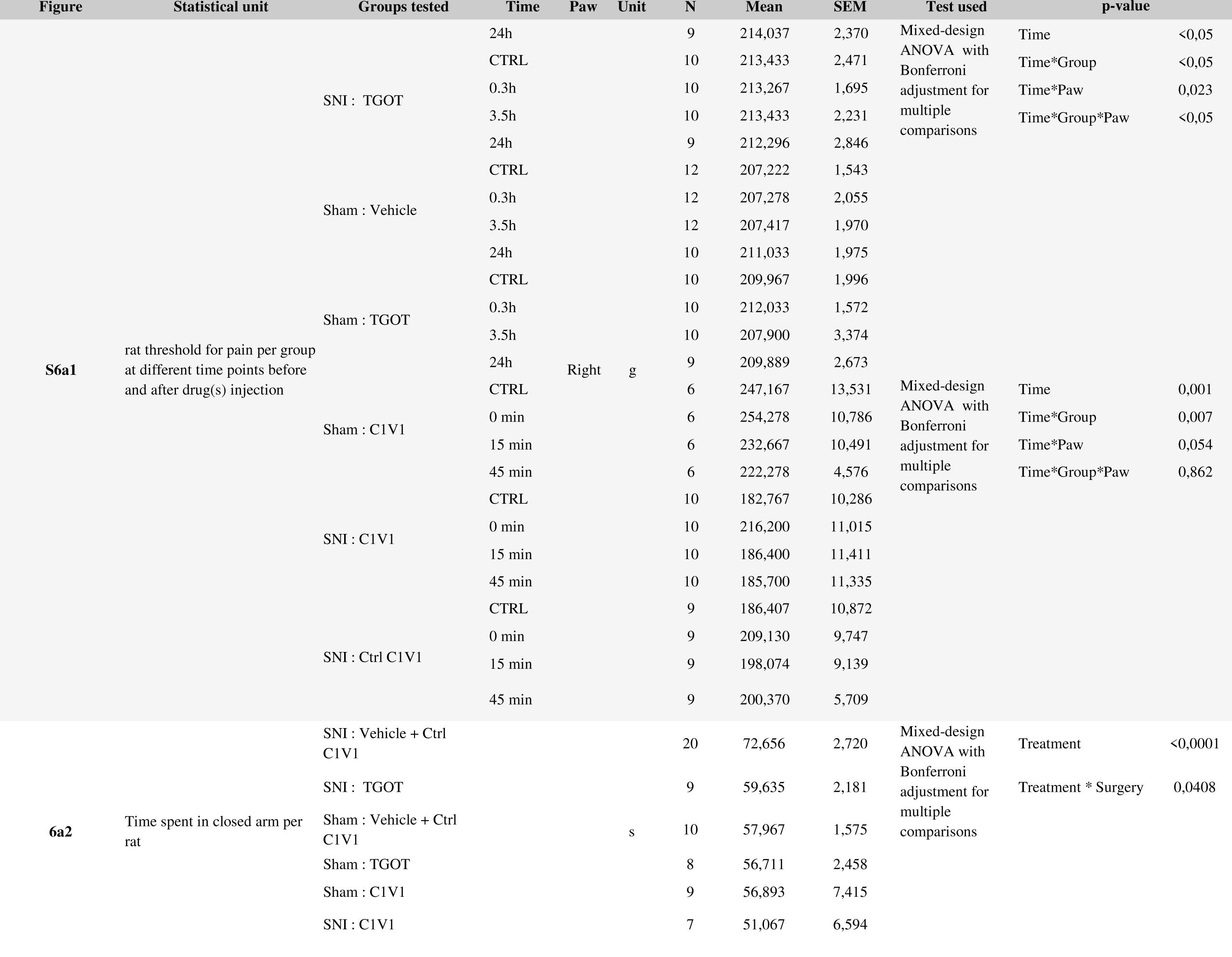
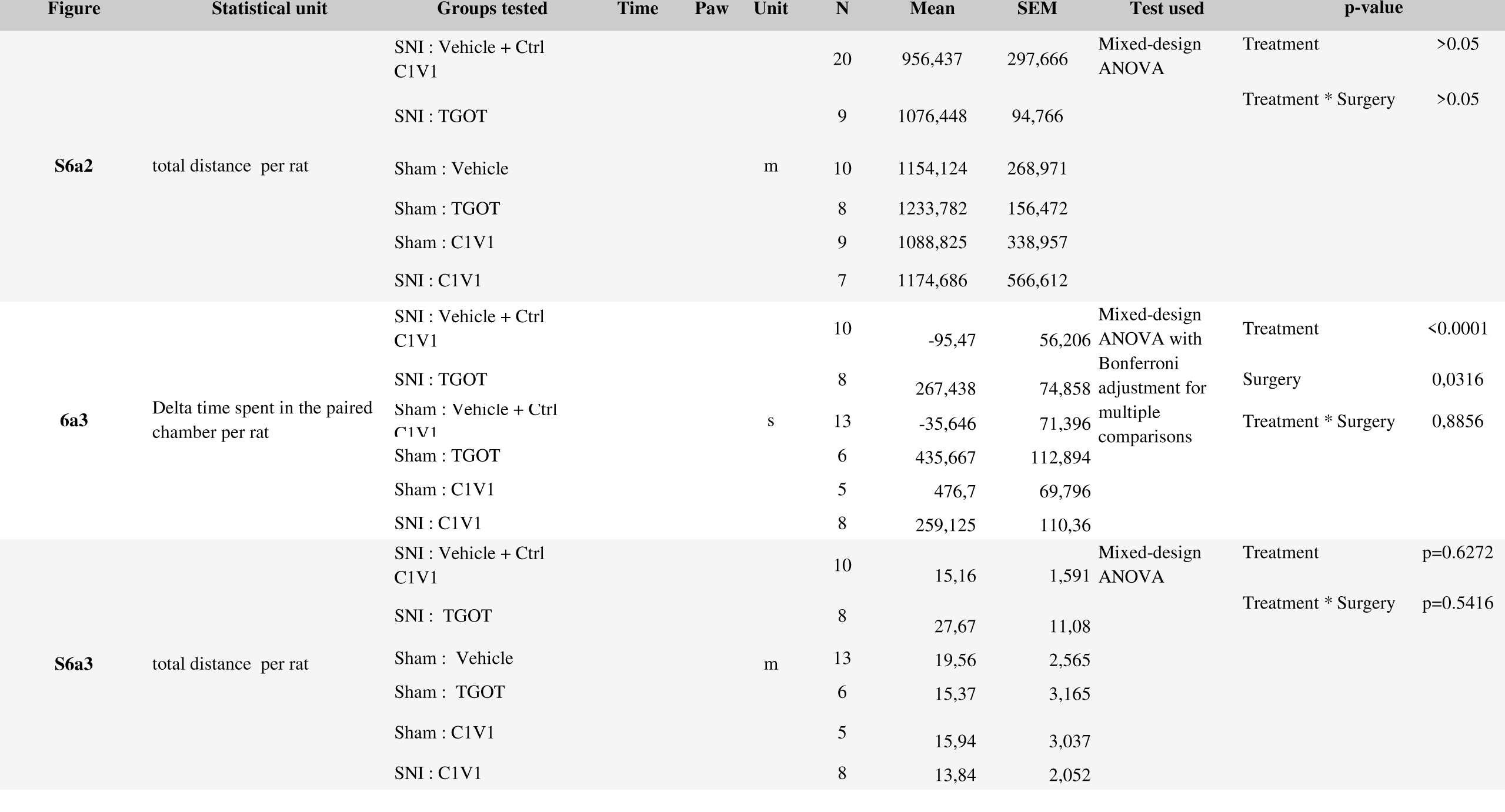
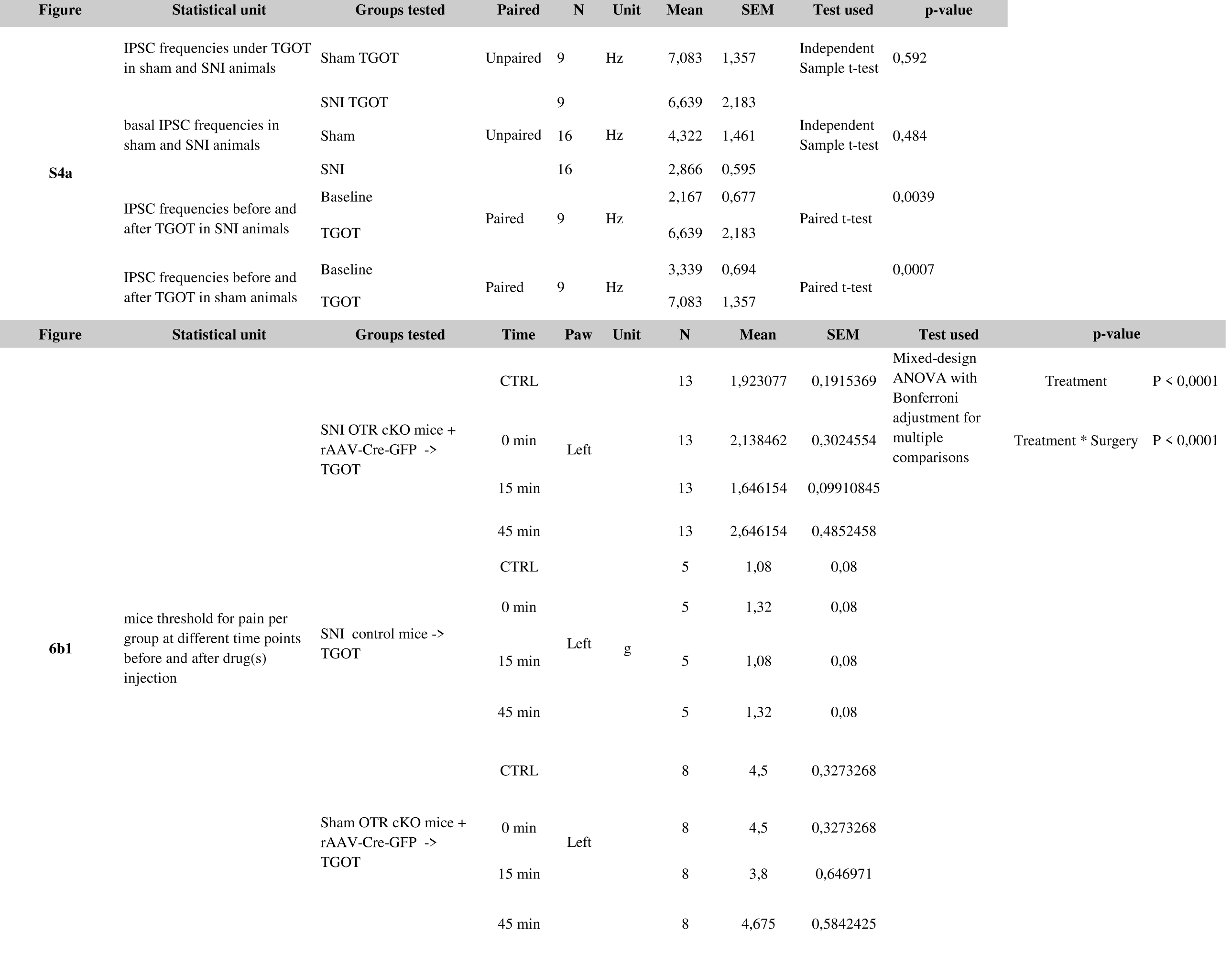
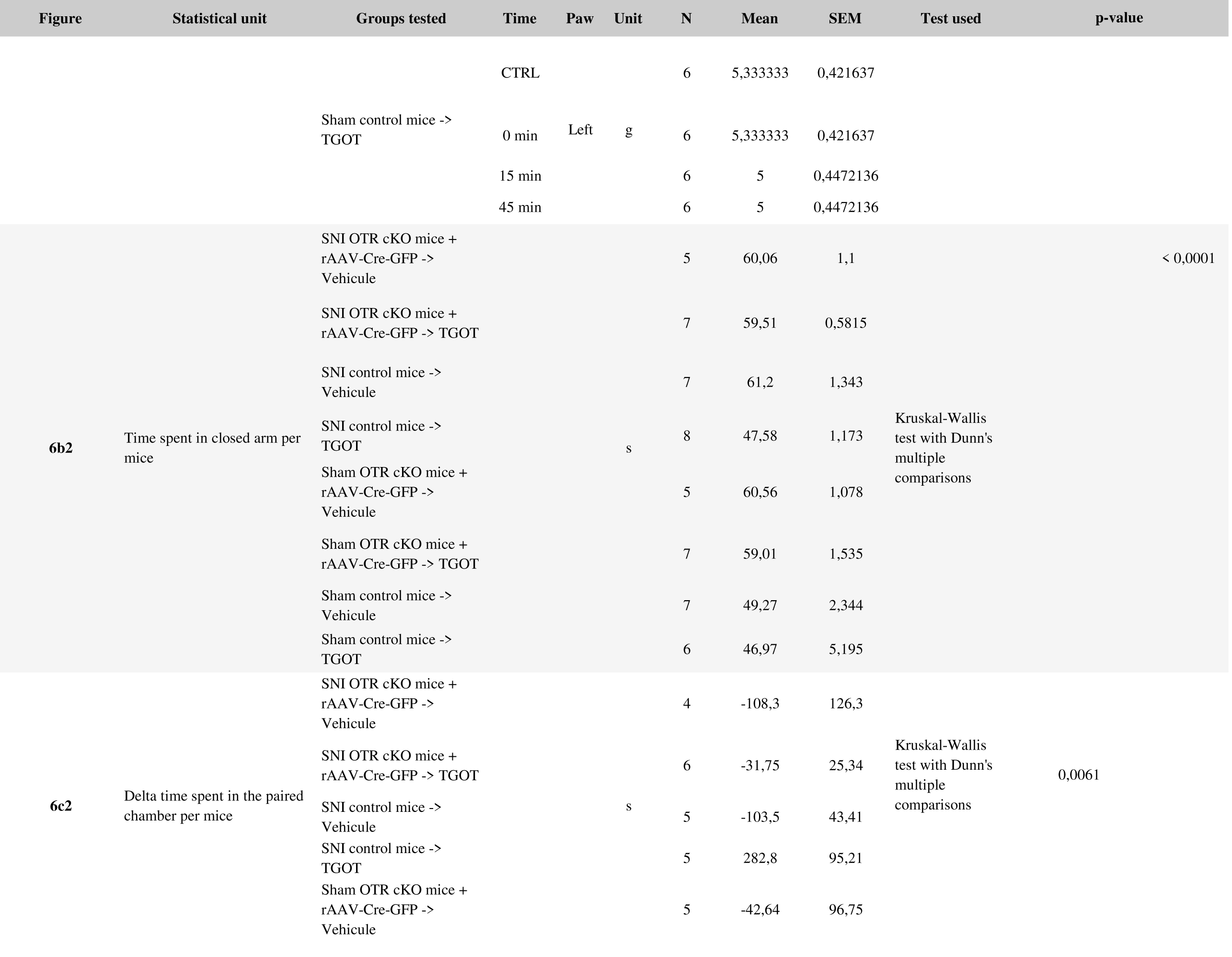
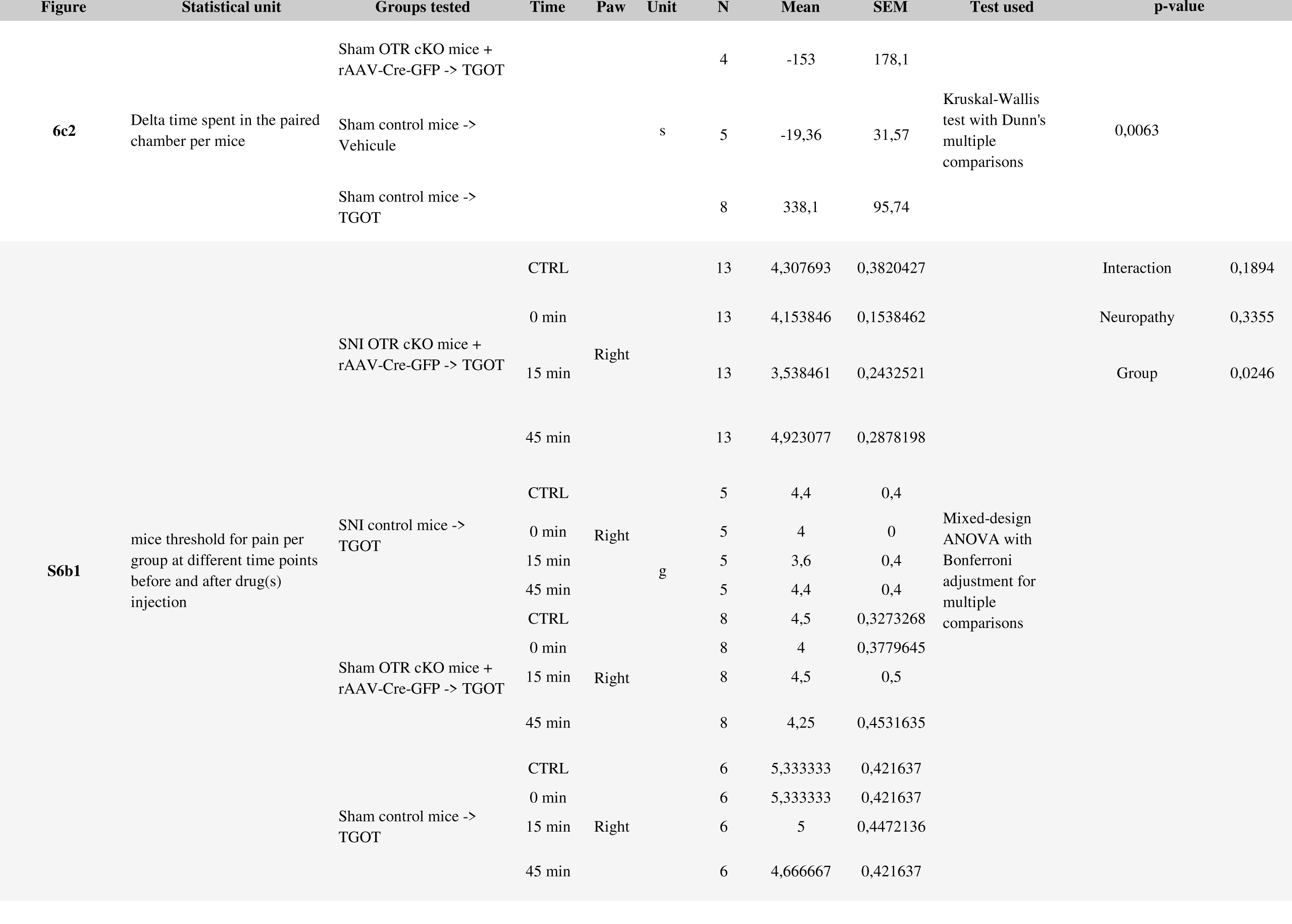
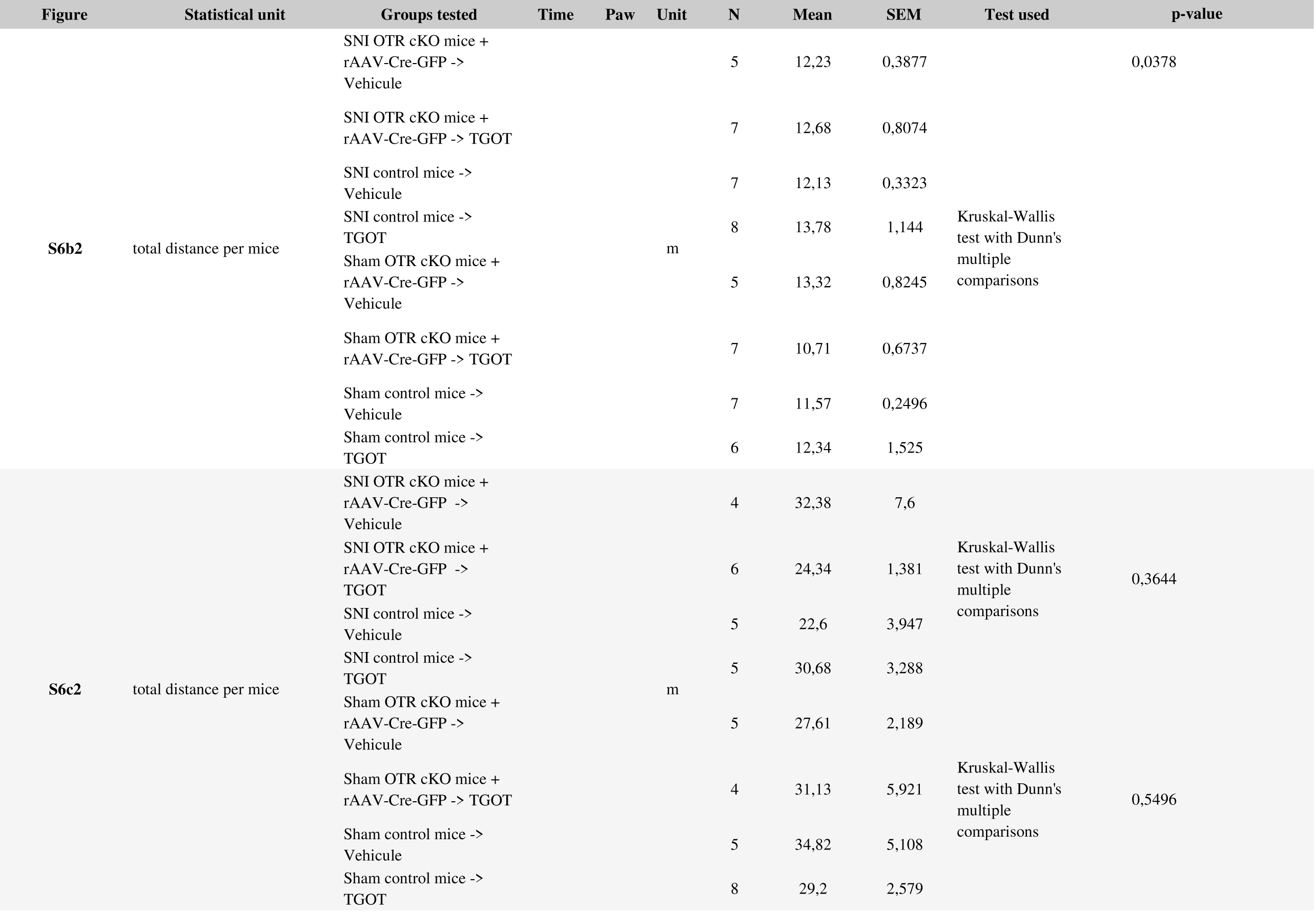
Numerical Values and Statistical Analysis of Data presented in Fig. 6 and **S6**

**Table S7.**
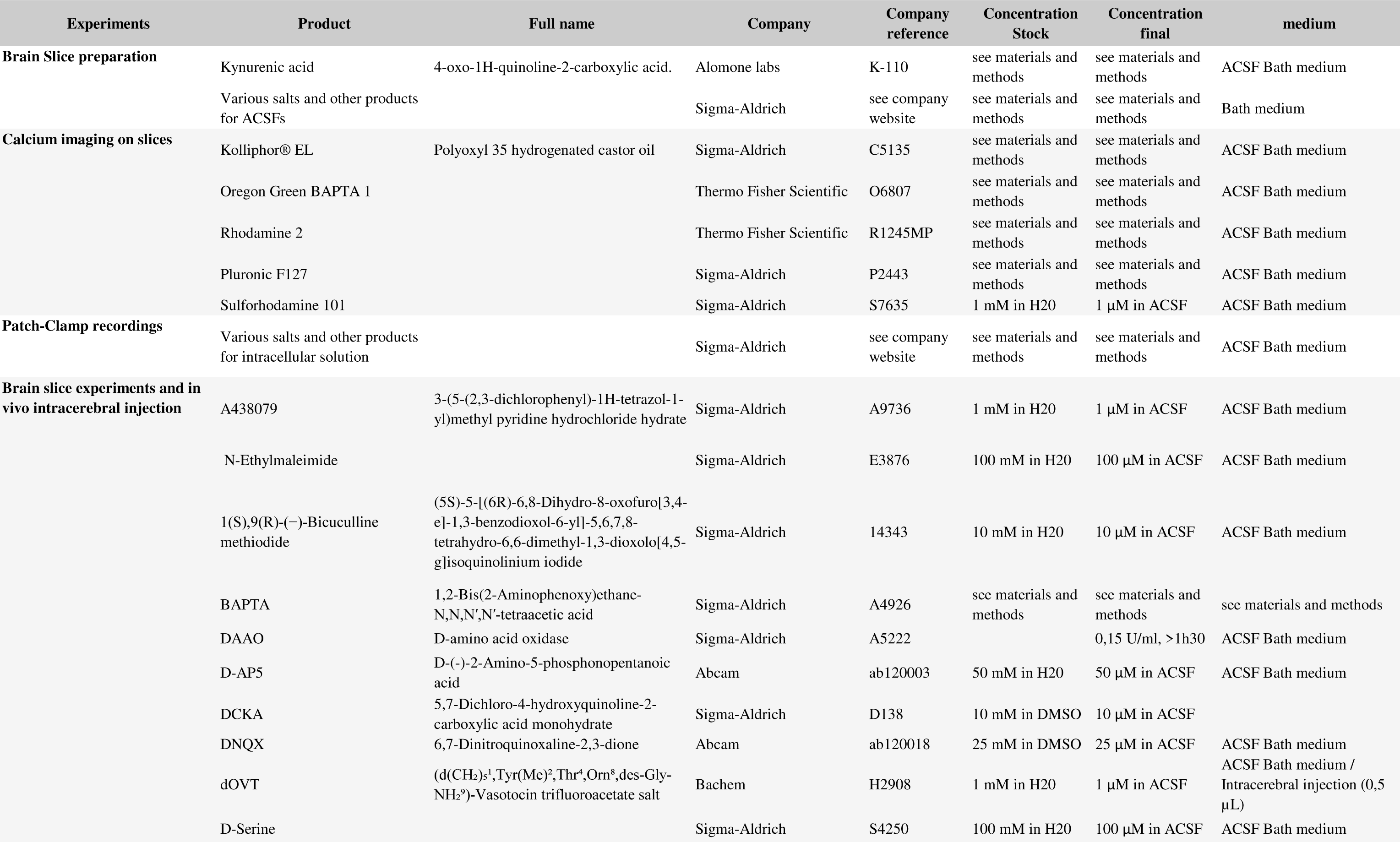
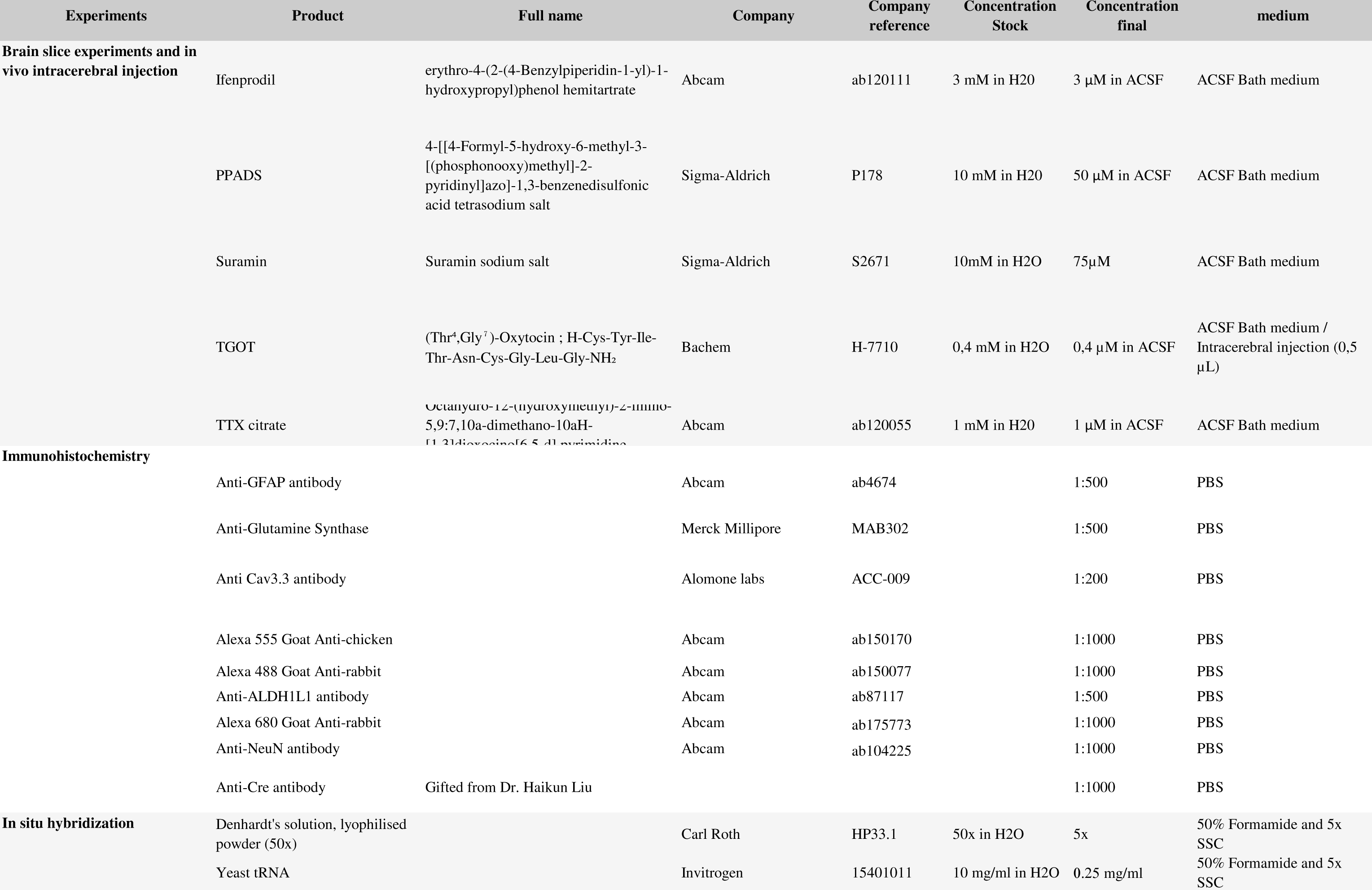
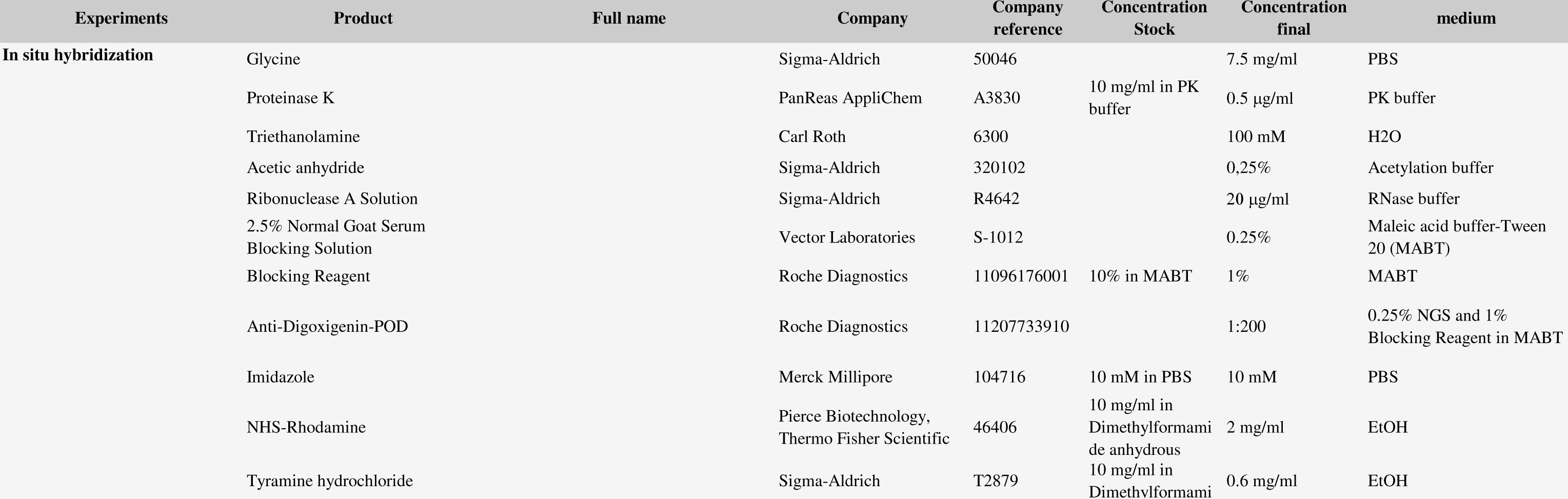
List of Reagents used in this Study

| Experiments | Product | Full name | Company | Company reference | Concentration Stock | Concentration final | medium |
| --- | --- | --- | --- | --- | --- | --- | --- |
| **Brain Slice preparation** | Kynurenic acid | 4-oxo-1H-quinoline-2-carboxylic acid. | Alomone labs | K-110 | see materials and methods | see materials and methods | ACSF Bath medium |
|  | Various salts and other products for ACSFs |  | Sigma-Aldrich | see company website | see materials and methods | see materials and methods | Bath medium |
| **Calcium imaging on slices** | Kolliphor® EL | Polyoxyl 35 hydrogenated castor oil | Sigma-Aldrich | C5135 | see materials and methods | see materials and methods | ACSF Bath medium |
|  | Oregon Green BAPTA 1 |  | Thermo Fisher Scientific | O6807 | see materials and methods | see materials and methods | ACSF Bath medium |
|  | Rhodamine 2 |  | Thermo Fisher Scientific | R1245MP | see materials and methods | see materials and methods | ACSF Bath medium |
|  | Pluronic F127 |  | Sigma-Aldrich | P2443 | see materials and methods | see materials and methods | ACSF Bath medium |
|  | Sulforhodamine 101 |  | Sigma-Aldrich | S7635 | 1 mM in H2O | 1 µM in ACSF | ACSF Bath medium |
| **Patch-Clamp recordings** | Various salts and other products for intracellular solution |  | Sigma-Aldrich | see company website | see materials and methods | see materials and methods | ACSF Bath medium |
| **Brain slice experiments and in vivo intracerebral injection** | A438079 | 3-(5-(2,3-dichlorophenyl)-1H-tetrazol-1-yl)methyl pyridine hydrochloride hydrate | Sigma-Aldrich | A9736 | 1 mM in H2O | 1 µM in ACSF | ACSF Bath medium |
|  | N-Ethylmaleimide |  | Sigma-Aldrich | E3876 | 100 mM in H2O | 100 µM in ACSF | ACSF Bath medium |
|  | 1(S),9(R)-(-)-Bicuculline methiodide | (5S)-5-[(6R)-6,8-Dihydro-8-oxofuro[3,4-e]-1,3-benzodioxol-6-yl]-5,6,7,8-tetrahydro-6,6-dimethyl-1,3-dioxolo[4,5-g]isoquinolinium iodide | Sigma-Aldrich | 14343 | 10 mM in H2O | 10 µM in ACSF | ACSF Bath medium |
|  | BAPTA | 1,2-Bis(2-Aminophenoxy)ethane-N,N,N',N'-tetraacetic acid | Sigma-Aldrich | A4926 | see materials and methods | see materials and methods | see materials and methods |
|  | DAAO | D-amino acid oxidase | Sigma-Aldrich | A5222 |  | 0,15 U/ml, >1h30 | ACSF Bath medium |
|  | D-AP5 | D-(-)-2-Amino-5-phosphonopentanoic acid | Abcam | ab120003 | 50 mM in H2O | 50 µM in ACSF | ACSF Bath medium |
|  | DCKA | 5,7-Dichloro-4-hydroxyquinoline-2-carboxylic acid monohydrate | Sigma-Aldrich | D138 | 10 mM in DMSO | 10 µM in ACSF |  |
|  | DNQX | 6,7-Dinitroquinoxaline-2,3-dione | Abcam | ab120018 | 25 mM in DMSO | 25 µM in ACSF | ACSF Bath medium |
|  | dOVT | (d(CH<sub>2</sub>)<sub>5</sub>)<sup>1</sup>,Tyr(Me)<sup>2</sup>,Thr<sup>4</sup>,Orn<sup>8</sup>,des-Gly-NH<sub>2</sub><sup>9</sup>)-Vasotocin trifluoroacetate salt | Bachem | H2908 | 1 mM in H2O | 1 µM in ACSF | ACSF Bath medium / Intracerebral injection (0,5 µL) |
|  | D-Serine |  | Sigma-Aldrich | S4250 | 100 mM in H2O | 100 µM in ACSF | ACSF Bath medium |
| Brain slice experiments and in vivo intracerebral injection | Ifenprodil | erythro-4-(2-(4-Benzylpiperidin-1-yl)-1-hydroxypropyl)phenol hemitartrate | Abcam | ab120111 | 3 mM in H2O | 3 µM in ACSF | ACSF Bath medium |
|  | PPADS | 4-[[[4-Formyl-5-hydroxy-6-methyl-3-[(phosphonoxy)methyl]-2-pyridinyl]azo]-1,3-benzenedisulfonic acid tetrasodium salt | Sigma-Aldrich | P178 | 10 mM in H2O | 50 µM in ACSF | ACSF Bath medium |
|  | Suramin | Suramin sodium salt | Sigma-Aldrich | S2671 | 10mM in H2O | 75µM | ACSF Bath medium |
|  | TGOT | (Thr<sup>4</sup>,Gly<sup>7</sup>)-Oxytocin ; H-Cys-Tyr-Ile-Thr-Asn-Cys-Gly-Leu-Gly-NH<sub>2</sub> | Bachem | H-7710 | 0,4 mM in H2O | 0,4 µM in ACSF | ACSF Bath medium / Intracerebral injection (0,5 µL) |
|  | TTX citrate | Octahydro-12-(hydroxymethyl)-2-imino-5,9:7,10a-dimethano-10aH-[1,3]dioxepino[6,5-d]pyrimidine | Abcam | ab120055 | 1 mM in H2O | 1 µM in ACSF | ACSF Bath medium |
| Immunohistochemistry |  |  |  |  |  |  |  |
|  | Anti-GFAP antibody |  | Abcam | ab4674 |  | 1:500 | PBS |
|  | Anti-Glutamine Synthase |  | Merck Millipore | MAB302 |  | 1:500 | PBS |
|  | Anti Cav3.3 antibody |  | Alomone labs | ACC-009 |  | 1:200 | PBS |
|  | Alexa 555 Goat Anti-chicken |  | Abcam | ab150170 |  | 1:1000 | PBS |
|  | Alexa 488 Goat Anti-rabbit |  | Abcam | ab150077 |  | 1:1000 | PBS |
|  | Anti-ALDH1L1 antibody |  | Abcam | ab87117 |  | 1:500 | PBS |
|  | Alexa 680 Goat Anti-rabbit |  | Abcam | ab175773 |  | 1:1000 | PBS |
|  | Anti-NeuN antibody |  | Abcam | ab104225 |  | 1:1000 | PBS |
|  | Anti-Cre antibody | Gifted from Dr. Haikun Liu |  |  |  | 1:1000 | PBS |
| In situ hybridization | Denhardt's solution, lyophilised powder (50x) |  | Carl Roth | HP33.1 | 50x in H2O | 5x | 50% Formamide and 5x SSC |
|  | Yeast tRNA |  | Invitrogen | 15401011 | 10 mg/ml in H2O | 0.25 mg/ml | 50% Formamide and 5x SSC |
| In situ hybridization | Glycine |  | Sigma-Aldrich | 50046 |  | 7.5 mg/ml | PBS |
|  | Proteinase K |  | PanReas AppliChem | A3830 | 10 mg/ml in PK buffer | 0.5 µg/ml | PK buffer |
|  | Triethanolamine |  | Carl Roth | 6300 |  | 100 mM | H2O |
|  | Acetic anhydride |  | Sigma-Aldrich | 320102 |  | 0,25% | Acetylation buffer |
|  | Ribonuclease A Solution |  | Sigma-Aldrich | R4642 |  | 20 µg/ml | RNase buffer |
|  | 2.5% Normal Goat Serum Blocking Solution |  | Vector Laboratories | S-1012 |  | 0.25% | Maleic acid buffer-Tween 20 (MABT) |
|  | Blocking Reagent |  | Roche Diagnostics | 11096176001 | 10% in MABT | 1% | MABT |
|  | Anti-Digoxigenin-POD |  | Roche Diagnostics | 11207733910 |  | 1:200 | 0.25% NGS and 1% Blocking Reagent in MABT |
|  | Imidazole |  | Merck Millipore | 104716 | 10 mM in PBS | 10 mM | PBS |
|  | NHS-Rhodamine |  | Pierce Biotechnology, Thermo Fisher Scientific | 46406 | 10 mg/ml in Dimethylformami de anhydrous | 2 mg/ml | EtOH |
|  | Tyramine hydrochloride |  | Sigma-Aldrich | T2879 | 10 mg/ml in Dimethylformami | 0.6 mg/ml | EtOH |

## REFERENCES

1. Lee, H. J., Macbeth, A. H., Pagani, J. H. & Scott Young, W. Oxytocin: The great facilitator of life. Prog. Neurobiol. 88, 127–151 (2009).

2. Eliava, M. et al. A New Population of Parvocellular Oxytocin Neurons Controlling Magnocellular Neuron Activity and Inflammatory Pain Processing. Neuron 89, 1291–304 (2016).

3. Huber, D., Veinante, P. & Stoop, R. Vasopressin and oxytocin excite distinct neuronal populations in the central amygdala. Science 308, 245–8 (2005).

4. Viviani, D. et al. Oxytocin selectively gates fear responses through distinct outputs from the central amygdala. Science 333, 104–7 (2011).

5. Knobloch, H. S. et al. Evoked axonal oxytocin release in the central amygdala attenuates fear response. Neuron 73, 553–66 (2012).

6. Hasan, M. T. et al. A Fear Memory Engram and Its Plasticity in the Hypothalamic Oxytocin System. Neuron 103, 133–146.e8 (2019).

7. Neugebauer, V., Li, W., Bird, G. C. & Han, J. S. The amygdala and persistent pain. Neuroscientist 10, 221–234 (2004).

8. Eto, K., Kim, S. K., Takeda, I. & Nabekura, J. The roles of cortical astrocytes in chronic pain and other brain pathologies. Neurosci. Res. 126, 3–8 (2018).

9. Hansen, R. R. & Malcangio, M. Astrocytes - Multitaskers in chronic pain. Eur. J. Pharmacol. 716, 120–128 (2013).

10. Ji, R.-R. R., Berta, T. & Nedergaard, M. Glia and pain: Is chronic pain a gliopathy? Pain 154, S10–S28 (2013).

11. Poisbeau, P., Grinevich, V. & Charlet, A. Oxytocin Signaling in Pain: Cellular, Circuit, System, and Behavioral Levels. Curr. Top. Behav. Neurosci. (2017). doi:10.1007/7854_2017_14

12. Rash, J. a, Aguirre-Camacho, A. & Campbell, T. S. Oxytocin and pain: a systematic review and synthesis of findings. Clin. J. Pain 30, 453–62 (2014).

13. Kuo, J., Hariri, O. R. & Micevych, P. An interaction of oxytocin receptors with metabotropic glutamate receptors in hypothalamic astrocytes. J. Neuroendocrinol. 21, 1001–6 (2009).

14. Parent, A.-S. et al. Oxytocin Facilitates Female Sexual Maturation through a Glia-to-Neuron Signaling Pathway. Endocrinology 149, 1358–1365 (2008).

15. Di Scala-Guenot, D., Mouginot, D. & Strosser, M.-T. Increase of intracellular calcium induced by oxytocin in hypothalamic cultures astrocytes. Glia 11, 269–276 (1994).

16. Wang, P., Qin, D. & Wang, Y.-F. Oxytocin Rapidly Changes Astrocytic GFAP Plasticity by Differentially Modulating the Expressions of pERK 1/2 and Protein Kinase A. Front. Mol. Neurosci. 10, 1–14 (2017).

17. Adamsky, A. et al. Astrocytic Activation Generates De Novo Neuronal Potentiation and Memory Enhancement. Cell 1–13 (2018). doi:10.1016/j.cell.2018.05.002

18. Allen, N. J. Astrocyte regulation of synaptic behavior. Annu. Rev. Cell Dev. Biol. 30, 439– 63 (2014).

19. Khakh, B. S. & Deneen, B. The Emerging Nature of Astrocyte Diversity. Annu. Rev. Neurosci. 42, 187–207 (2019).

20. Verkhratsky, A. & Nedergaard, M. Physiology of Astroglia. Physiol. Rev. 98, 239–389 (2018).

21. Martin-Fernandez, M. et al. Synapse-specific astrocyte gating of amygdala-related behavior. Nat. Neurosci. 20, 1540–1548 (2017).

22. Bazargani, N. & Attwell, D. Astrocyte calcium signaling: the third wave. Nat. Neurosci. 19, 182–189 (2016).

23. Savtchouk, I. & Volterra, A. Gliotransmission: Beyond Black-and-White. J. Neurosci. 38, 14–25 (2018).

24. Shigetomi, E., Patel, S. & Khakh, B. S. Probing the Complexities of Astrocyte Calcium Signaling. Trends Cell Biol. 26, 300–12 (2016).

25. Lee, H. J., Caldwell, H. K., Macbeth, A. H., Tolu, S. G. & Young, W. S. A conditional knockout mouse line of the oxytocin receptor. Endocrinology 149, 3256–3263 (2008).

26. Perea, G. & Araque, A. Properties of synaptically evoked astrocyte calcium signal reveal synaptic information processing by astrocytes. J. Neurosci. 25, 2192–203 (2005).

27. Shigetomi, E., Bowser, D. N., Sofroniew, M. V & Khakh, B. S. Two forms of astrocyte calcium excitability have distinct effects on NMDA receptor-mediated slow inward currents in pyramidal neurons. J. Neurosci. 28, 6659–63 (2008).

28. Jourdain, P. et al. Glutamate exocytosis from astrocytes controls synaptic strength. Nat. Neurosci. 10, 331–9 (2007).

29. Mothet, J.-P. et al. Glutamate receptor activation triggers a calcium-dependent and SNARE protein-dependent release of the gliotransmitter D-serine. Proc. Natl. Acad. Sci. 102, 5606–5611 (2005).

30. Oliet, S. H. R. & Mothet, J.-P. Regulation of N-methyl-D-aspartate receptors by astrocytic D-serine. Neuroscience 158, 275–83 (2009).

31. Robin, L. M. et al. Astroglial CB1 Receptors Determine Synaptic D-Serine Availability to Enable Recognition Memory. Neuron 0, 1–10 (2018).

32. Neugebauer, V., Galhardo, V., Maione, S. & Mackey, S. C. Forebrain pain mechanisms. Brain Res. Rev. 60, 226–42 (2009).

33. Sah, P., Faber, E. S. L., Lopez De Armentia, M. & Power, J. The amygdaloid complex: anatomy and physiology. Physiol. Rev. 83, 803–34 (2003).

34. Alba-Delgado, C., Cebada-Aleu, A., Mico, J. A. & Berrocoso, E. Comorbid anxiety-like behavior and locus coeruleus impairment in diabetic peripheral neuropathy: A comparative study with the chronic constriction injury model. Prog. Neuropsychopharmacol. Biol. Psychiatry 71, 45–56 (2016).

35. Yalcin, I. et al. The sciatic nerve cuffing model of neuropathic pain in mice. J. Vis. Exp. 1– 7 (2014). doi:10.3791/51608

36. Han, J. S. & Neugebauer, V. Synaptic plasticity in the amygdala in a visceral pain model in rats. Neuroscience Letters 361, (2004).

37. Phelps, E. A. & LeDoux, J. E. Contributions of the Amygdala to Emotion Processing: From Animal Models to Human Behavior. Neuron 48, 175–187 (2005).

38. Yalcin, I. & Barrot, M. The anxiodepressive comorbidity in chronic pain. Curr Opin Anaesthesiol 27, 520–527 (2014).

39. Grinevich, V. & Stoop, R. Interplay between Oxytocin and Sensory Systems in the Orchestration of Socio-Emotional Behaviors. Neuron 99, 887–904 (2018).

40. Ferretti, V. et al. Oxytocin Signaling in the Central Amygdala Modulates Emotion Discrimination in Mice. Curr. Biol. 29, 1938–1953.e6 (2019).

41. Hung, L. W. et al. Gating of social reward by oxytocin in the ventral tegmental area. Science 357, 1406–1411 (2017).

42. Marlin, B. J., Mitre, M., D’amour, J. A., Chao, M. V. & Froemke, R. C. Oxytocin enables maternal behaviour by balancing cortical inhibition. Nature 520, 499–504 (2015).

43. Menon, R. et al. Oxytocin Signaling in the Lateral Septum Prevents Social Fear during Lactation. Curr. Biol. 28, 1066–1078.e6 (2018).

44. Nardou, R. et al. Oxytocin-dependent reopening of a social reward learning critical period with MDMA. Nature 569, 116–120 (2019).

45. Oettl, L. L. et al. Oxytocin Enhances Social Recognition by Modulating Cortical Control of Early Olfactory Processing. Neuron 90, 609–621 (2016).

46. Xiao, L., Priest, M. F., Nasenbeny, J., Lu, T. & Kozorovitskiy, Y. Biased Oxytocinergic Modulation of Midbrain Dopamine Systems. Neuron 95, 368–384.e5 (2017).

47. Mitre, M. et al. A Distributed Network for Social Cognition Enriched for Oxytocin Receptors. J. Neurosci. 36, 2517–2535 (2016).

48. Yoshida, M. et al. Evidence that oxytocin exerts anxiolytic effects via oxytocin receptor expressed in serotonergic neurons in mice. J. Neurosci. 29, 2259–71 (2009).

49. Fellin, T. et al. Neuronal synchrony mediated by astrocytic glutamate through activation of extrasynaptic NMDA receptors. Neuron 43, 729–43 (2004).

50. Angulo, M. C., Kozlov, A. S., Charpak, S. & Audinat, E. Glutamate released from glial cells synchronizes neuronal activity in the hippocampus. J. Neurosci. 24, 6920–6927 (2004).

51. Durkee, C. A. et al. G i/o protein-coupled receptors inhibit neurons but activate astrocytes and stimulate gliotransmission. Glia 67, 1076–1093 (2019).

52. Busnelli, M. et al. Functional selective oxytocin-derived agonists discriminate between individual G protein family subtypes. J. Biol. Chem. 287, 3617–3629 (2012).

53. Chini, B., Verhage, M. & Grinevich, V. The Action Radius of Oxytocin Release in the Mammalian CNS: From Single Vesicles to Behavior. Trends Pharmacol. Sci. 38, 982–991 (2017).

54. Grinevich, V., Knobloch-Bollmann, H. S., Eliava, M., Busnelli, M. & Chini, B. Assembling the Puzzle: Pathways of Oxytocin Signaling in the Brain. Biol. Psychiatry 79, 155–164 (2016).

55. Hirase, H., Iwai, Y., Takata, N., Shinohara, Y. & Mishima, T. Volume transmission signalling via astrocytes. Philos. Trans. R. Soc. B Biol. Sci. 369, 20130604–20130604 (2014).

56. Kastanenka, K. V. et al. A roadmap to integrate astrocytes into Systems Neuroscience. GLIA (2019). doi:10.1002/glia.23632

57. Oliveira, J. F., Sardinha, V. M., Guerra-Gomes, S., Araque, A. & Sousa, N. Do stars govern our actions? Astrocyte involvement in rodent behavior. Trends Neurosci. 38, 535– 549 (2015).

58. Volterra, A., Liaudet, N. & Savtchouk, I. Astrocyte Ca(2+) signalling: an unexpected complexity. Nat. Rev. Neurosci. 15, 327–35 (2014).

59. Haim, L. Ben & Rowitch, D. Functional diversity of astrocytes in neural circuit regulation. Nat. Rev. Neurosci. 18, 31–41 (2016).

60. Khakh, B. S. & Sofroniew, M. V. Diversity of astrocyte functions and phenotypes in neural circuits. Nat. Neurosci. 18, 942–52 (2015).

61. Filous, A. R. & Silver, J. “Targeting astrocytes in CNS injury and disease: A translational research approach”. Prog. Neurobiol. 144, 173–187 (2016).

62. Marlin, B. J. & Froemke, R. C. Oxytocin modulation of neural circuits for social behavior. Dev. Neurobiol. 77, 169–189 (2017).

63. Morales-Soto, W. & Gulbransen, B. D. Enteric Glia: A New Player in Abdominal Pain. Cell. Mol. Gastroenterol. Hepatol. 7, 433–445 (2019).

64. Poisbeau, P., Grinevich, V. & Charlet, A. Oxytocin Signaling in Pain: Cellular, Circuit, System, and Behavioral Levels. in Current Topics in Behavioral Neurosciences 289–320 (Springer, Berlin, Heidelberg, 2017). doi:10.1007/7854

65. Bender, C. L., Calfa, G. D. & Molina, V. A. Astrocyte plasticity induced by emotional stress: A new partner in psychiatric physiopathology? Prog. Neuro-Psychopharmacology Biol. Psychiatry 65, 68–77 (2016).

66. Meyer-Lindenberg, A., Domes, G., Kirsch, P. & Heinrichs, M. Oxytocin and vasopressin in the human brain: social neuropeptides for translational medicine. Nat. Rev. Neurosci. 12, 524–38 (2011).

67. Shigetomi, E. et al. Imaging calcium microdomains within entire astrocyte territories and endfeet with GCaMPs expressed using adeno-associated viruses. J. Gen. Physiol. 141, 633–647 (2013).

68. Decosterd, I. & Woolf, C. J. Spared nerve injury: an animal model of persistent peripheral neuropathic pain. Pain 87, 149–158 (2000).

69. Ikegaya, Y., Le Bon-Jego, M. & Yuste, R. Large-scale imaging of cortical network activity with calcium indicators. Neurosci. Res. 52, 132–138 (2005).

70. Serrano, A., Haddjeri, N., Lacaille, J., Robitaille, R. & Centre-ville, S. GABAergic Network Activation of Glial Cells Underlies Hippocampal Heterosynaptic Depression. J. Neurosci. 26, 5370–5382 (2006).

71. Anlauf, E. & Derouiche, A. Glutamine synthetase as an astrocytic marker: its cell type and vesicle localization. Front. Endocrinol. (Lausanne). 4, 144 (2013).

72. Yizhar, O. et al. Neocortical excitation/inhibition balance in information processing and social dysfunction. Nature 477, 171–178 (2011).

73. Luis-Delgado, O. E. et al. Calibrated forceps: A sensitive and reliable tool for pain and analgesia studies. J. Pain 7, 32–39 (2006).

74. Walf, A. A. & Frye, C. A. The use of the elevated plus maze as an assay of anxiety-related behavior in rodents. Nat. Protoc. 2, 322–8 (2007).

75. King, T. et al. Unmasking the tonic-aversive state in neuropathic pain. Nat. Neurosci. 12, 1364–1366 (2009).

76. Clark, P. J. & Evans, F. C. Distance to Nearest Neighbor as a Measure of Spatial Relationships in Populations. Ecology 35, 445–453 (1954).

77. Millet, L. J., Collens, M. B., Perry, G. L. W. & Bashir, R. Pattern analysis and spatial distribution of neurons in culture. Integr. Biol. 3, 1167 (2011).

